# On the validity of electric brain signal predictions based on population firing rates

**DOI:** 10.1101/2024.07.10.602833

**Authors:** Torbjørn V. Ness, Tom Tetzlaff, Gaute T. Einevoll, David Dahmen

## Abstract

Neural activity at the population level is commonly studied experimentally through measurements of electric brain signals like local field potentials (LFPs), or electroencephalography (EEG) signals. To allow for comparison between observed and simulated neural activity it is therefore important that simulations of neural activity can accurately predict these brain signals. Simulations of neural activity at the population level often rely on point-neuron network models or firing-rate models. While these simplified representations of neural activity are computationally efficient, they lack the explicit spatial information needed for calculating LFP/EEG signals. Different heuristic approaches have been suggested for overcoming this limitation, but the accuracy of these approaches has not fully been assessed. One such heuristic approach, the so-called kernel method, has previously been applied with promising results and has the additional advantage of being well-grounded in the biophysics underlying electric brain signal generation. It is based on calculating rate-to-LFP/EEG kernels for each synaptic pathway in a network model, after which LFP/EEG signals can be obtained directly from population firing rates. This amounts to a massive reduction in the computational effort of calculating brain signals because the brain signals are calculated for each population instead of for each neuron. Here, we investigate how and when the kernel method can be expected to work, and present a theoretical framework for predicting its accuracy. We show that the relative error of the brain signal predictions is a function of the single-cell kernel heterogeneity and the spike-train correlations. Finally, we demonstrate that the kernel method is most accurate for the dominating brain signal contributions. We thereby further establish the kernel method as a promising approach for calculating electric brain signals from large-scale neural simulations.

## 1. Introduction

Science is at its most productive when models can make experimental predictions so that experimental results can inform and improve the models. Measurable brain signals should therefore be available from simulations of neural activity. The brain is studied at many different scales, from the molecular scale to behavior, and the different scales rely on models at different levels of abstraction. It is therefore important to have well-founded methods to calculate different types of brain signals, from neural simulations at different levels of abstraction (Figure 1) [Einevoll et al., 2019].

**Figure 1:**
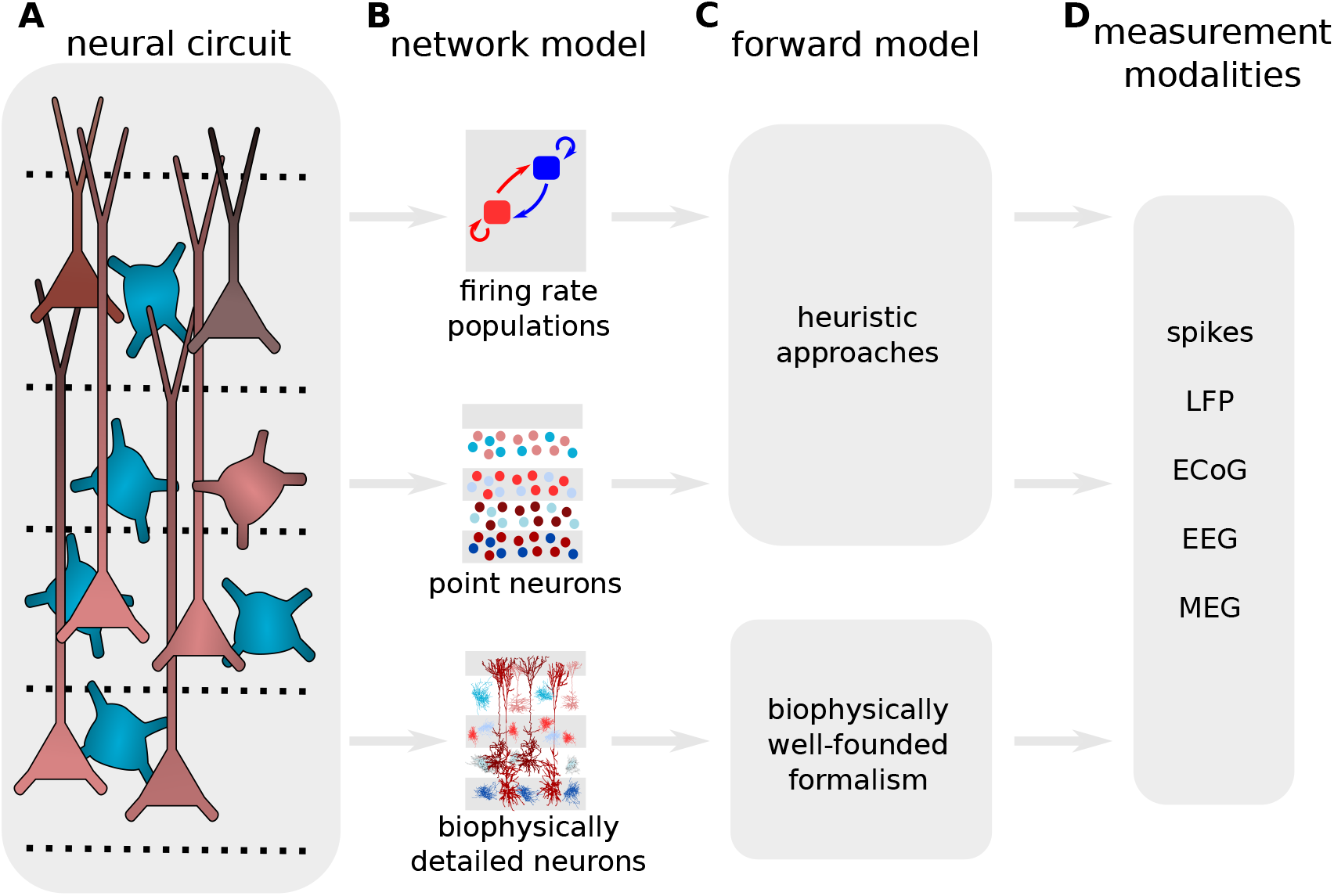
Measurable signals should be available from neural simulations at different levels of abstraction. Neural circuits, here represented by a putative cortical column (panel **A**), are studied at different levels of biological detail, depending on the scientific question (panel **B**). By using a forward model (panel **C**) one can calculate measurable signals (panel **D**) from neural activity simulated at different levels of abstraction. In general, calculations of such brain signals are only biophysically well-founded when using biophysically detailed cell models, while simplified representations of neurons will require “heuristic” approaches where it can be hard to estimate the accuracy of the resulting brain signal predictions.

To study neural activity at the level of neural populations, it is common to rely on measurements of local field potentials (LFPs), which is the low-frequency part of the extracellular potential measured inside the brain, or on electroencephalography (EEG) signals, which is the extracellular potential measured outside of the head. The most accurate way to calculate LFP and EEG signals from simulated neural network activity is to use biophysically detailed multicompartment neuron models, coupled with volume conductor theory [Holt and Koch, 1999; Einevoll et al., 2013; Ness et al., 2022; Halnes et al., 2024]. For single neurons or small populations, this is in principle straight-forward [Hagen et al., 2018; Næss et al., 2021], and this approach has been pursued also for large recurrently connected networks by a handful of studies [Reimann et al., 2013; Tomsett et al., 2015; Hagen et al., 2018; Dai et al., 2020; Baratham et al., 2022; Borges et al., 2022; Rimehaug et al., 2023; Romani et al., 2024].

However, biophysically detailed modeling of neural activity at the population levels is extremely computationally demanding, and often not viable in practice [Einevoll et al., 2019]. Therefore, when studying neural network activity, it is more common to rely on simplified representations of neurons and neural activity, through for example point-neuron network models [Gerstner et al., 2014; Potjans and Diesmann, 2014; Billeh et al., 2020] or firing-rate models [Deco et al., 2008; Gerstner et al., 2014; Sanz-Leon et al., 2015]. These simplified representations of neural activity are more computationally tractable and typically orders of magnitude faster than biophysically detailed simulations [Billeh et al., 2020], but many brain signals, like the LFP and EEG signals, are generated by spatially distributed neural membrane currents, which are not available from the simplified schemes (since the spatial structure of individual neurons is not explicitly modeled) [Halnes et al., 2024]. An important topic is therefore what is the best approach for calculating approximations of different brain signals from neural activity simulated from point-neuron network simulations or firing-rate models.

Several different approaches to calculate LFP/EEG/MEG signals from point-neuron or firing-rate models have been suggested [Deco et al., 2008; Sanz-Leon et al., 2015; Mazzoni et al., 2015; Hagen et al., 2016; Teleńczuk et al., 2020a; Martínez-Cañada et al., 2021; Glomb et al., 2022; Tesler et al., 2022], but quantitative evaluations of the accuracy of such approaches have often been hard to come by, due to the lack of “ground truth” signals to compare the approximations to. It has therefore often been unclear how well these approximations work, although there are important exceptions that we discuss later.

A common approach with a long history to get approximate LFP/EEG signals from firing-rate models is simply to assume that the signal is proportional to the firing rate [Deco et al., 2008]. For EEG/MEG signals, it is sometimes instead assumed that an equivalent current dipole is proportional to the firing rate, and the dipole can be inserted into a head model to obtain the EEG/MEG signal [Sanz-Leon et al., 2015]. Although this approach can certainly be useful, it neglects some basic principles in how these signals are generated, and as a result, some error will be introduced in the time-domain of the predicted signals [Mazzoni et al., 2015; Hagen et al., 2022; Halnes et al., 2024].

Hagen et al. [2016] presented the so-called “hybrid scheme”, where the neural network activity is first simulated in a point-neuron network, and saved to file. Afterward, in a separate step, the spiking activity is replayed onto biophysically detailed cell models, from which the resulting LFP signals can be calculated. The hybrid scheme is a computationally expensive approach because it relies on representing all neurons that are within the reach of the recording electrode [Lindén et al., 2011; Kajikawa and Schroeder, 2011] with a high level of morphological and electrophysiological detail. On the other hand, it is well grounded in the biophysics of extracellular signal generation.

Hagen et al. [2016] also used the hybrid scheme to test a “kernel approach”, where they calculated LFP kernels for each synaptic pathway in the model. Each population kernel represented the average postsynaptic LFP contribution given an action potential in the presynaptic population, and the LFP signal could then be approximated by convolving the firing rate of each presynaptic population with the corresponding population kernel and summing the LFP contributions for each synaptic pathway in the model. This kernel approach was confirmed to give accurate approximations to the LFP, at a very low computational cost once the kernels were known because the LFP could be predicted directly from the firing rate of each population, instead of from the transmembrane currents of each individual neuron. A major drawback of this approach was that the calculation of the population kernels was still very computationally demanding.

Mazzoni et al. [2015] tested so-called “proxy” methods for calculating LFP signals (later also extended to EEG signals by Martínez-Cañada et al. [2021]) directly from point-neuron network simulations, and found that a weighted sum of synaptic currents, which are available from point-neuron network simulations, could be used to predict the LFP calculated by a more comprehensive approach, similar to the “hybrid scheme” discussed above. The proxies were demonstrated to be quite accurate and provided excellent LFP predictions for the use-case considered. On the other hand, they are in a sense phenomenological and typically poorly grounded in the underlying biophysics of extracellular potentials, which can in some cases be a drawback.

Teleńczuk et al. [2020a] used experimentally measured LFP kernels from spike-trigger averaged LFP recordings, and used these kernels to approximate LFP signals, by convolving them with firing rates from point-neuron network simulations. This approach has the advantage of being independent of the modeling choices that are required when simulating LFP kernels [Teleńczuk et al., 2020a,b]. This approach was later expanded upon by Tesler et al. [2022], to also enable MEG signal predictions from point-neuron network models or firing-rate models. However, kernels measured from spike-triggered averages are potentially troubled by correlations, and Hagen et al. [2016] obtained different results when calculating kernels directly, and from spike-triggered averages, even within the same model. This can also be directly observed, as the measured kernels are not always causal, which we would expect them to be given that they represent the postsynaptic contribution from a presynaptic spike. Further, the measured excitatory LFP kernels were proposed to be disynaptic inhibitory kernels [Teleńczuk et al., 2017; Teleńczuk et al., 2020a], illustrating a problem with interpreting results based on LFP kernels from spike-triggered averages. Note that the degree to which measured spike-triggered LFP kernels are contaminated by correlations will depend on the scenario. For example for the monosynaptic thalamic activation of cortical postsynaptic target cells considered by Swadlow et al. [2002], the contamination was very small.

The earlier attempts to model LFP kernels have required a large number of single-cell simulations [Hagen et al., 2016, 2017; Teleńczuk et al., 2020b] to represent the postsynaptic population. However, a very efficient yet highly biophysically detailed framework for calculating population kernels was recently proposed by Hagen et al. [2022]. In this framework, a single biophysically detailed cell simulation was sufficient to accurately predict a population kernel by first obtaining the membrane currents of the single postsynaptic neuron in response to conductance-based synaptic input, and letting this represent the population-averaged membrane currents following synaptic activation. All other effects, including the spatial extent of the population and the variability of synaptic parameters, were then accounted for by a series of linear convolutions in the spatial and temporal domains. This approach greatly increases the applicability of the kernel approach, since LFP/EEG kernels can be calculated accurately and efficiently, even by common laptop computers. The LFP calculated from the kernel approach by Hagen et al. [2022] was tested against the “ground truth” LFP calculated from a multicompartment, biophysically detailed neural network simulation, and the kernel approach was found to be quite accurate in most scenarios.

As reviewed above, several recent projects have used the kernel approach to estimate LFP, EEG, or MEG signals directly from firing rates [Hagen et al., 2016; Teleńczuk et al., 2020a,b; Skaar et al., 2020; Hagen et al., 2022; Tesler et al., 2022], and it has proved a promising tool for future studies of neural activity at the population level. Therefore, it is important to have a good qualitative understanding of how the kernel approach works, and good quantitative measures of how accurate it is under different circumstances.

In this study, we start by building a better understanding of how and when the kernel method can be expected to work, and when caution is advised. We then develop a theoretical framework for predicting the accuracy of the kernel approach and show that the relative error is a function of the single-cell kernel heterogeneity and spike-train correlations. Finally, we demonstrate that the kernel approach is most accurate for the LFP contributions that can be expected to dominate the LFP signal, like highly concentrated and correlated synaptic input to large populations of pyramidal neurons.

## 2. Results

Many measurable brain signals, like LFPs, ECoGs, EEGs, and MEGs are expected to share the same biophysical origin, namely the membrane currents following large numbers of synaptic inputs to populations of geometrically aligned pyramidal neurons [Ness et al., 2022; Halnes et al., 2024]. To accurately calculate these signals from simulated neural activity, we therefore need to take into account all synaptic events.

Since volume conduction is linear [Miceli et al., 2017], the compound extracellular potential

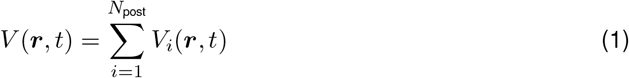

generated by a population of *N*_post_ neurons is a linear superposition of the individual cell contributions *V*_*i*_ (*i* = 1, …, *N*_post_). Therefore, calculating the extracellular potential of a population of *N*_post_ neurons is typically done by focusing on the synaptic input to each cell, calculating single-cell contributions (see Methods), and finally summing all cells (Figure 2**A**), here referred to as the *postsynaptic perspective*.

**Figure 2:**
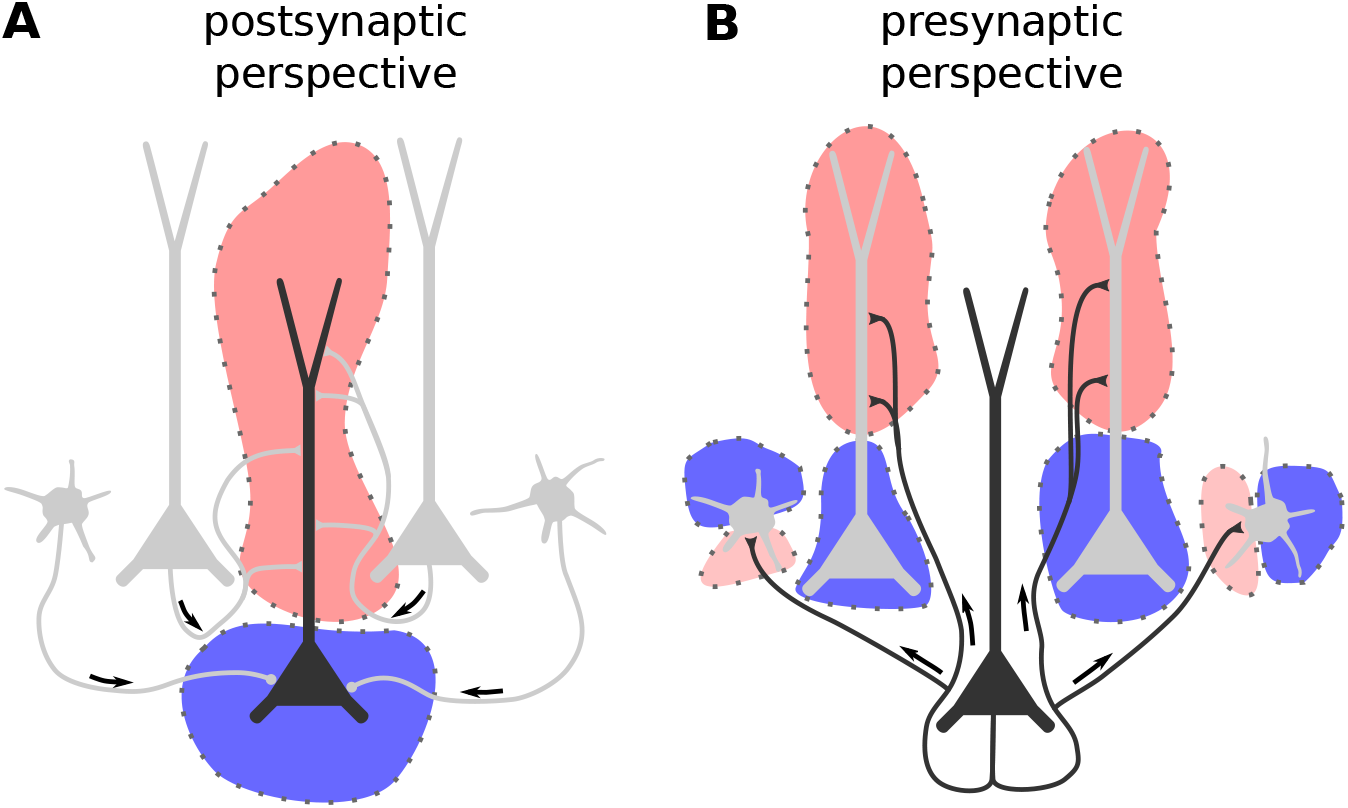
Illustration of the principle underlying the kernel method. **A:** The postsynaptic perspective, where all incoming synaptic input to a postsynaptic cell is taken into account, and the time-dependent LFP contribution of the postsynaptic cell is calculated. The total population LFP *V* (***r***, *t*) is then the sum of all such single-cell contributions *V*_*i*_(***r***, *t*). This is the standard way of calculating LFP signals from neural simulations. **B:** The presynaptic perspective, where all outgoing synapses from a single cell are considered. For passive cells with static (no plasticity), current-based synapses, every action potential of a presynaptic neuron *j* will evoke the same postsynaptic currents, and hence, each action potential has a fixed LFP response *h*_*ij*_ (***r***, *t*). By taking into account all postsynaptic targets, the single-cell kernel *k*_*j*_ (***r***, *t*) can be calculated, and the single-cell LFP contribution can be found by convolving the single-cell kernel with the corresponding spike train of the presynaptic cell. The population LFP is again the sum of all single-cell contributions, and if this is done for all cells, and all external incoming synapses, the LFP calculated by these two approaches will be identical, under the assumptions listed above.

### 2.1. Single-cell spike-LFP kernels

In principle, we can also switch the perspective to each presynaptic cell: Each action potential from a given cell leads to an activation of the outgoing synapses, causing a distributed “extracellular potential flash” from all postsynaptic target cells (Figure 2**B**), referred to as the *single-cell spike-LFP kernel*. For simplicity, we will here refer to this as the single-cell kernel. If we convolve the single-cell kernel with the spike train of the presynaptic neuron, we get the extracellular potential including all postsynaptic effects from this neuron. If we know all single-cell kernels and corresponding spike trains, we can then calculate the extracellular potential as the sum of all single-cell postsynaptic contributions. If this is also done for external input, we have accounted for all synaptic events.

The above argument is based on the fact that the single-cell kernel is similar each time a neuron spikes. This holds if we ignore synaptic plasticity and assume that extracellular potential contributions caused by individual synaptic activations superimpose linearly for each cell. However, in principle, the membrane currents of a cell depend on the joint effect of all its spiking inputs, for example, active dendritic channels or voltage-dependent synaptic currents cause nonlinear interference of inputs. However, previous work has shown that LFPs can be well predicted with quasi-linear approximations of ion channels [Ness et al., 2016, 2018], and that kernel-based approaches can give accurate LFP predictions also for conductance-based synapses [Hagen et al., 2022]. In this case, the above assumption holds and we obtain

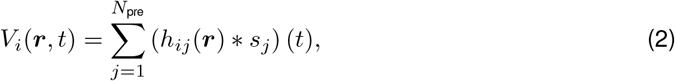

where *h*_*ij*_(***r***, *t*) is the LFP response of postsynaptic neuron *i* to an individual spike of presynaptic neuron *j*, and *s*_*j*_ is the spike train of presynaptic neuron *j*. Here * denotes a temporal convolution. If we combine equation (1) and equation (2) and rearrange summands, then we get what we refer to as the *presynaptic perspective* (Figure 2**B**),

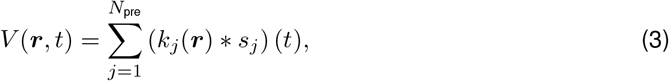

with the single-cell kernel

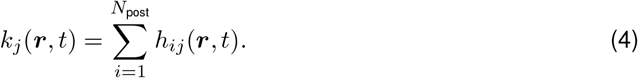

This prediction of the population LFP from single-cell kernels is in the following denoted as the “ground truth” against which we test approximations.

### 2.2. Population rate-LFP kernels

Neurons in neural circuits often share statistical properties in terms of morphology, electro-physiology, connections, and spiking activity. Based on such similarities they can be grouped into neuronal populations. In the classical view, a population is a group of neurons with similar input statistics as well as similar internal properties and dynamics, such that they have similar spiking statistics. For the generation of LFP contributions, however, not only the spiking statistics should be similar for cells within a population, but also their translation into LFPs as measured by the single-cell kernels.

If all single-cell kernels *k*_*j*_ of a population of neurons were identical, then they would in particular be identical to the population-averaged kernel

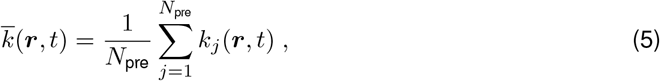

such that the compound LFP 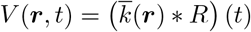 of the population could be perfectly predicted by the population rate 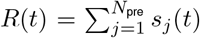 without the need to consider the detailed information of individual neuronal spike trains. The population-averaged kernel 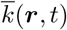 can therefore be interpreted as a *population rate-LFP kernel*. For simplicity, we will here refer to this as the population kernel.

In general, however, the properties and projections of neurons are only statistically similar rather than identical, such that the single-cell kernels differ from 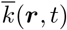. As a consequence

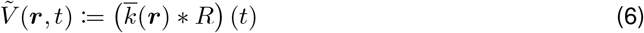

is only an approximation of the true compound LFP *V*(***r***, *t*). In the following, we study the error of this approximation and how it depends on the neuronal and the network properties.

Single-cell kernels depend on a multitude of neuron and network features including network connectivity, neuronal morphology, synapse positions, electrode position and electrical properties of cells, leading to potentially complicated spatio-temporal profiles. Yet they are by definition causal and their time course is determined by synaptic dynamics and dendritic filtering properties [Lindén et al., 2010]. As with LFP responses to individual synaptic inputs, the amplitude and polarity of single-cell kernels is expected to strongly depend on the relative position of cells with respect to the recording electrodes.

Before calculating the precise shape of single-cell kernels from biophysically detailed models, we first show some key aspects of the population kernel approximation using a simple illustrative model, where single-cell kernels are defined as double-exponential functions with different amplitudes (see Methods, Figure 3A).

**Figure 3:**
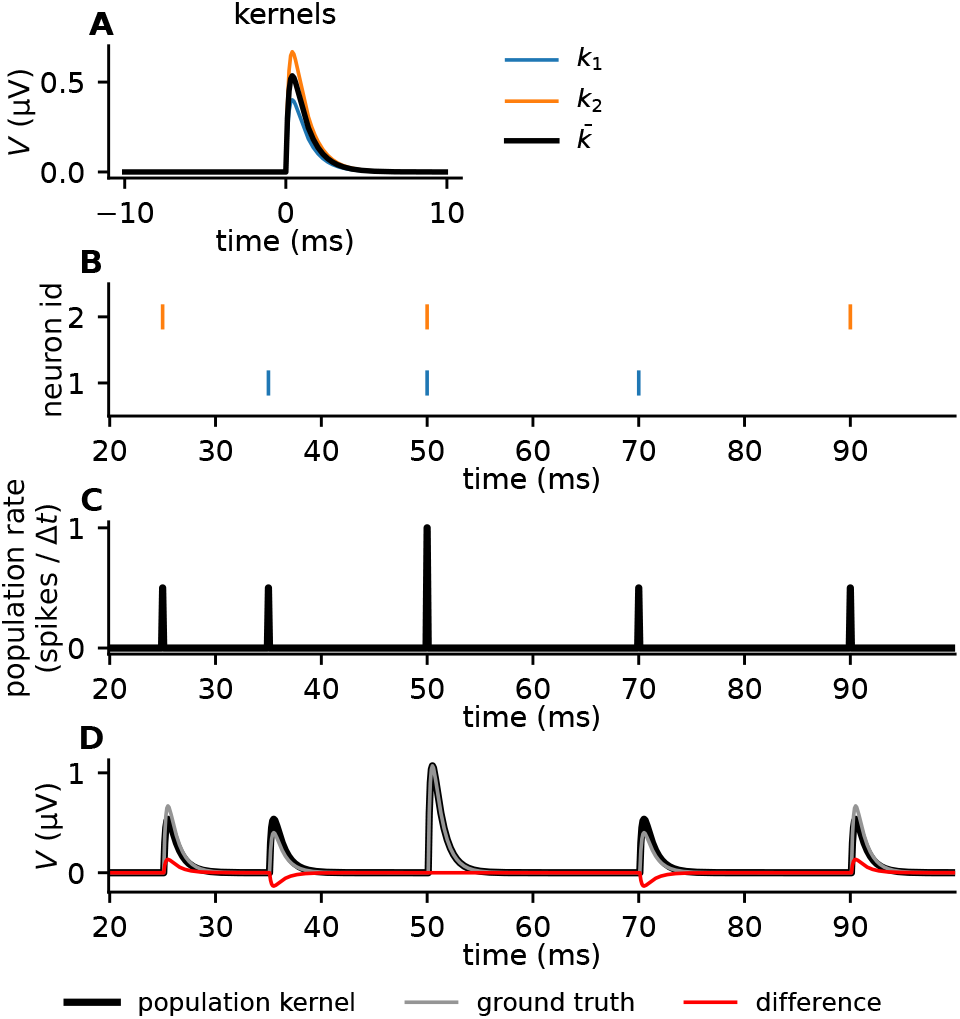
Illustration of kernel method with toy model. **A:** Two toy single-cell kernels (blue and orange), and the mean, that is, the population kernel (black). **B:** Raster plot of the two corresponding spike trains, with the same color code as in panel A. Each colored marker corresponds to a spike, and the individual spike trains are plotted at different heights along the *y*-axis. **C:** The population rate (average number of spikes per time bin, Δ*t* = 0.1 ms), that is, the mean firing rate from the spike trains in panel B. **D:** The gray line shows the ground truth toy LFP signal calculated as the sum of each single-cell contribution, which is again calculated by convolving the single-cell kernels with the corresponding spike trains. The black line shows the LFP calculated by convolving the population kernel with the population rate. The red line shows the difference between the ground truth LFP and the population kernel LFP.

Each single-cell kernel (Figure 3A) is convolved with a different spike train (Figure 3B) and the resulting extracellular potential (Figure 3D) is compared to the prediction of the population kernel (black line in Figure 3A) that is convolved with the population rate (Figure 3C). The population kernel prediction generally resembles the ground truth. It is, however, different in detail due to the heterogeneity in single-cell spike kernels. The approximation improves at times where multiple neurons spike synchronously (Figure 3D *t*=50 ms). This hints at a more general aspect: if all spike trains in equation (1) are identical, then the population kernel prediction becomes exact even though the single-cell kernels are different. In conclusion, this simple toy model illustrates the two main features that determine the quality of the population kernel prediction: spike-kernel heterogeneity and spike-train correlations. Predictions become poor when spike-train correlations are low and spike-kernel heterogeneity is large, whereas large spike-train correlations and low spike-kernel heterogeneity lead to low errors (Figure 4).

**Figure 4:**
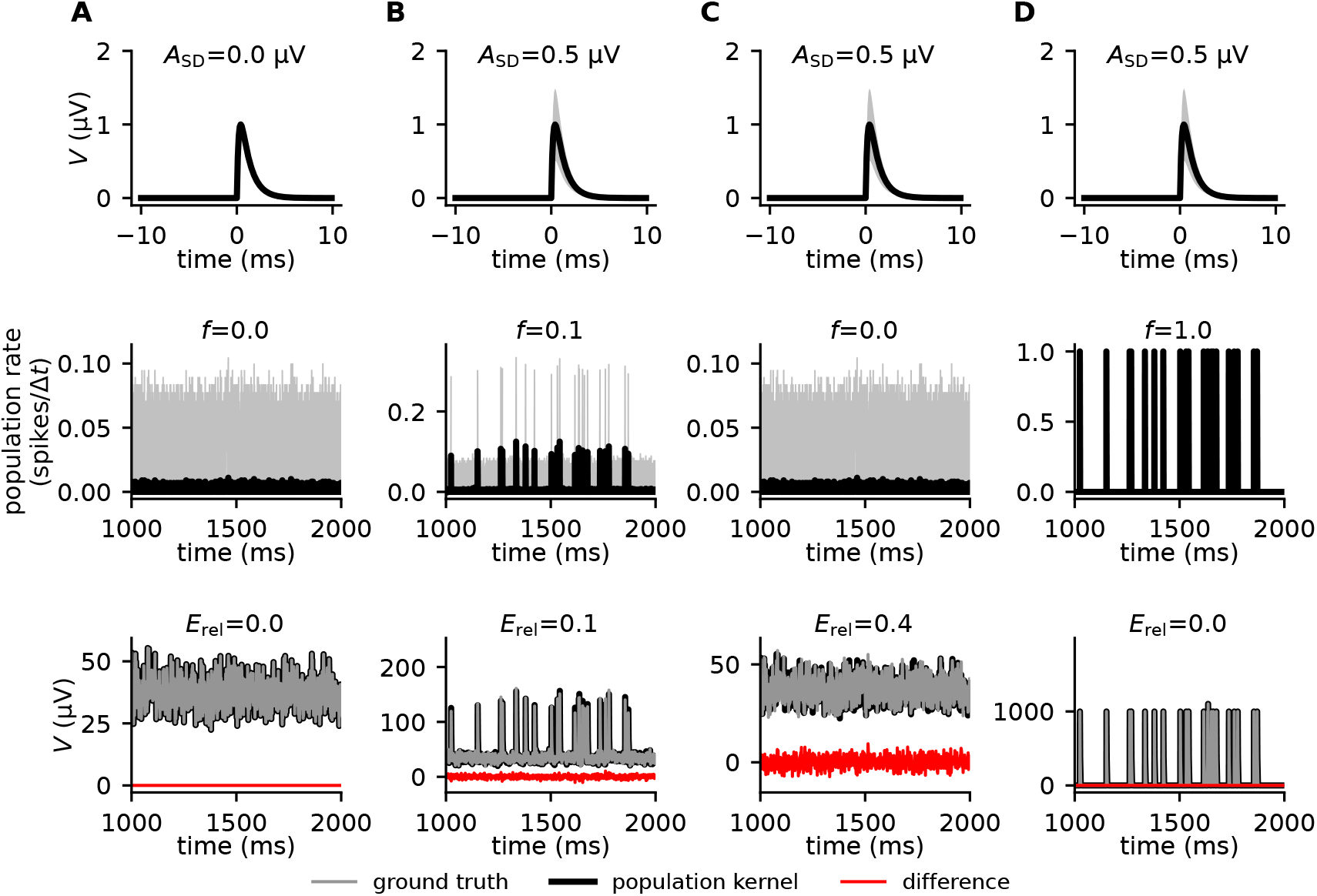
Error in population kernel predictions depend on kernel heterogeneity and spike-train correlations. Each column shows 1000 single-cell kernels with different amplitude standard deviations *A*_SD_ (top), and different levels of spike-train correlations (middle). Spike trains with varying levels of correlations were generated through Multiple Interaction Processes (MIP) [Kuhn et al., 2003], controlled by the parameter *f*, where *f* = 0 corresponds to uncorrelated homogeneous Poisson processes, while *f* = 1 corresponds to fully correlated (identical) spike trains (see Methods). The mean firing rate is shown in black, and the standard deviation in gray. The toy LFP is calculated (bottom). Relative error *E*_rel_, quantified by the normalized standard deviation of the difference between the ground truth signal and the population kernel signal (see Methods), vanishes for identical kernels, regardless of correlation (first column). For variable kernels with some correlation, the kernel approach will result in some relative error (second column). For variable kernels and zero correlation, the relative error will be large (third column). For perfect correlation, the relative error vanishes regardless of kernel variability (fourth column).

This behavior of the prediction error can be derived analytically by employing a statistical description of the setup. As mentioned above, a population of neurons is defined via statistical similarities between neuronal spike trains and spike kernels. In the following, we assume that both quantities, appearing as a product (convolution) in equation (3), are drawn from distributions with known means and covariances. A natural first choice for the definition of the prediction error would be the mean deviation Mean 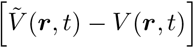 of the population kernel prediction 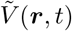 from the ground truth *V* (***r***, *t*), where Mean [·] denotes the average across time. We could then ask what this quantity is on expectation across different realizations of kernels. In fact, it is zero, because each individual single-cell kernel on expectation coincides with the expectation of the population-averaged kernel. This measure is therefore not informative about the prediction error of the population kernel method for a single realization of single-cell kernels. The error is better assessed by the standard deviation discrepancy of the population kernel prediction from the ground truth. The squared error then is 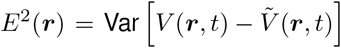, where Var [·] denotes the variance across time. The expectation of this quantity can be computed analytically (see Methods)

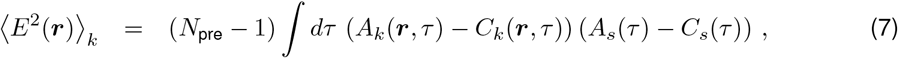

with ⟨·⟩_*k*_ denoting the expectation across realizations of the kernels. We further introduced the population averaged spike-train autocovariances *A*_*s*_(*τ*), the population averaged spike-train cross-covariances *C*_*s*_(*τ*), and the autocorrelation and cross-correlation of single-cell kernels *A*_*k*_(***r***, *τ*) = ∫ *dτ′* ⟨*k*_*i*_(***r***, *τ′*)*k*_*i*_(***r***, *τ′* + *τ*)⟩_*k*_, *C*_*k*_(***r***, *τ*) = ∫ *dτ′* ⟨*k*_*i*_(***r***, *τ′*)*k*_*j*_(***r***, *τ′* + *τ*)⟩. The expression for *E*^2^ shows that, as expected, the error vanishes if the population of neurons spikes in a fully correlated manner (*C*_*s*_ = *A*_*s*_) or if all neurons have the same spike-LFP kernels (*A*_*k*_ = *C*_*k*_). For low average cross-covariances *C*_*s*_ ≈ 0 as observed in cortex, the error is primarily determined by the size of the presynaptic population *N*_pre_, i.e., the number of single-cell kernels, the correlations in spike-LFP kernels, and the spike-train autocovariances. To assess the overall performance of the population-based prediction, it is useful to also consider the relative error *E*_rel_, defined as the expected error *E* normalized by the standard deviation of the ground-truth signal (see Methods). For our toy model, the analytical predictions for the absolute and the relative error perfectly match the results of numerical simulations (Figure 5). Theory and simulation confirm the anticipated trend that the error grows with increasing kernel heterogeneity and decreasing spike-train correlations (Figure 5B,D). The effect of spike-train correlations is, however, much more pronounced in the relative error (Figure 5C,E), as can be explained by the theory (see Appendix B). Note that the relative error is low in regions where the signal amplitude is large (Figure 5A).

**Figure 5:**
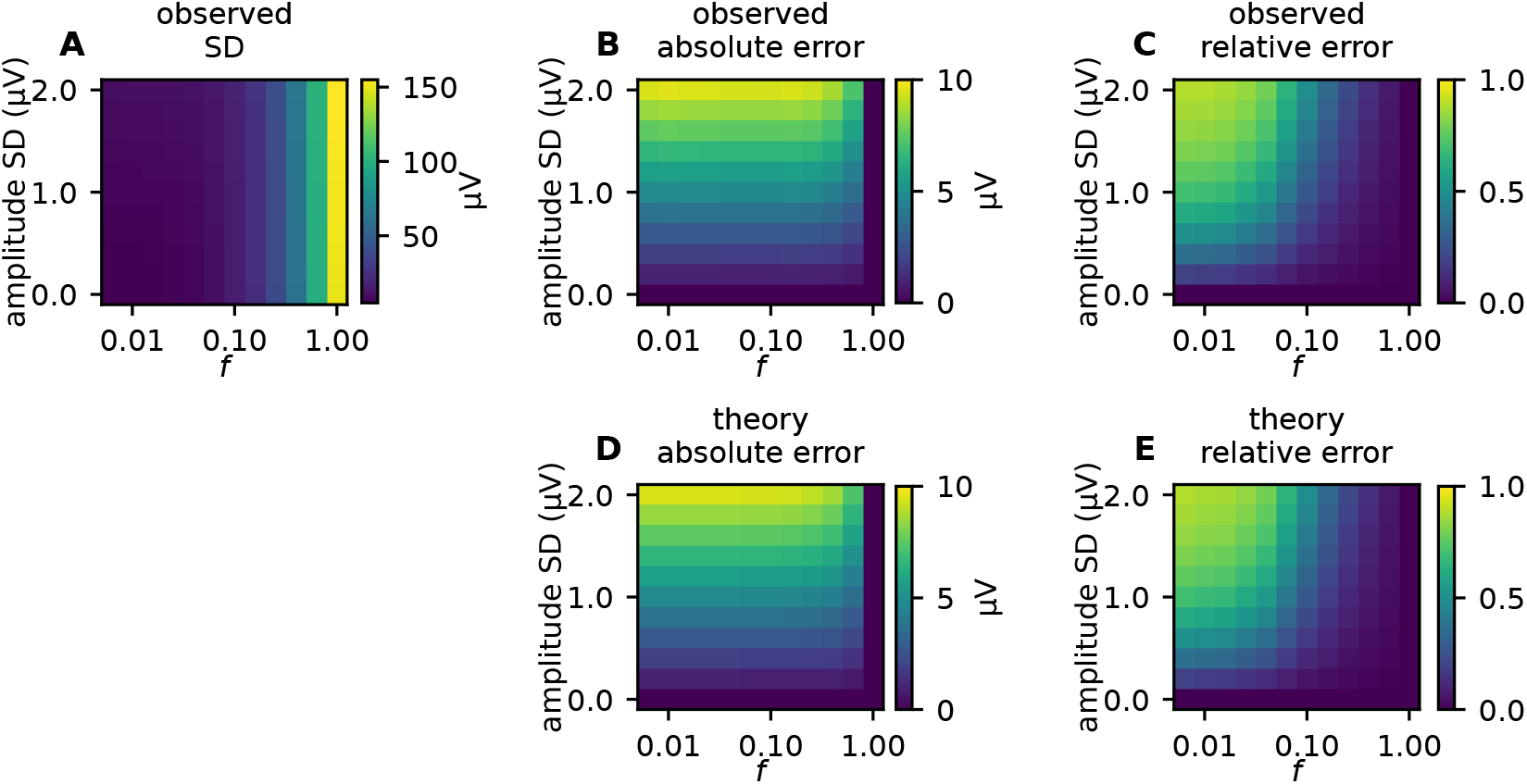
Parameter scan for simple toy-model LFP. **A**: The LFP amplitude (quantified by its standard deviation across time) for different levels of amplitude variability in single-cell kernels, and different levels of correlations between spike trains. **B**: Observed absolute error, quantified by the standard deviation of the difference between the ground truth signal and the population kernel signal. **C**: Observed relative error, quantified by the standard deviation of the difference between the ground truth signal and the population kernel signal (panel B), normalized by the ground truth signal amplitude (panel A). **D, E**: Same as in panels B and C, but predicted from theory (equation (7)). Correlated spike trains were generated using MIP processes (see Methods).

### 2.3. Sources and effect of kernel heterogeneities

As we have seen, the error depends on single-cell kernel heterogeneities. After having derived the general dependence of the population kernel prediction on the statistics of single-cell kernels and spike-train correlations, we next investigate more systematically where heterogeneity in single-cell kernels stems from. To this end, we need to go beyond the toy model of the previous section and employ a mechanistic model of extracellular potential generation based on the spatial distribution of cells, connectivity specifications and biophysically detailed cell models.

We consider LFP and EEG signals from cortical populations. The major contribution to these signals stems from synaptic inputs onto pyramidal neurons [Hagen et al., 2016; Halnes et al., 2024]. In the following, we therefore investigate the LFP and EEG kernels of a population of layer 5 pyramidal neurons, positioned around a linear multi-contact electrode that records the LFP at different depths, while the EEG is recorded outside the scalp (Figure 6A). Synaptic inputs from a single presynaptic neuron are modeled as spikes delivered to a random subset of neurons in the considered postsynaptic population. To account for the natural heterogeneity in cortical connectivity, parameters such as synapse locations, synaptic strengths, time constants, and delays are randomly drawn from predefined distributions. The calculations of postsynaptic membrane currents and resulting extracellular potentials are based on a morphologically reconstructed pyramidal neuron from Hay et al. [2011]. A single-cell kernel represents the post-synaptic LFP (EEG) response to the firing of a single presynaptic neuron. The population kernel corresponds to the average of the single-cell kernels obtained for different presynaptic neurons, each targeting different subsets of neurons in the postsynaptic population. Each population kernel represents one specific synaptic pathway from a given presynaptic population to a given postsynaptic population. Details of the setup outlined here are described in Figure 6 and Methods. In the following, we assess the sources of kernel heterogeneities by systematically varying the different features of this setup.

**Figure 6:**
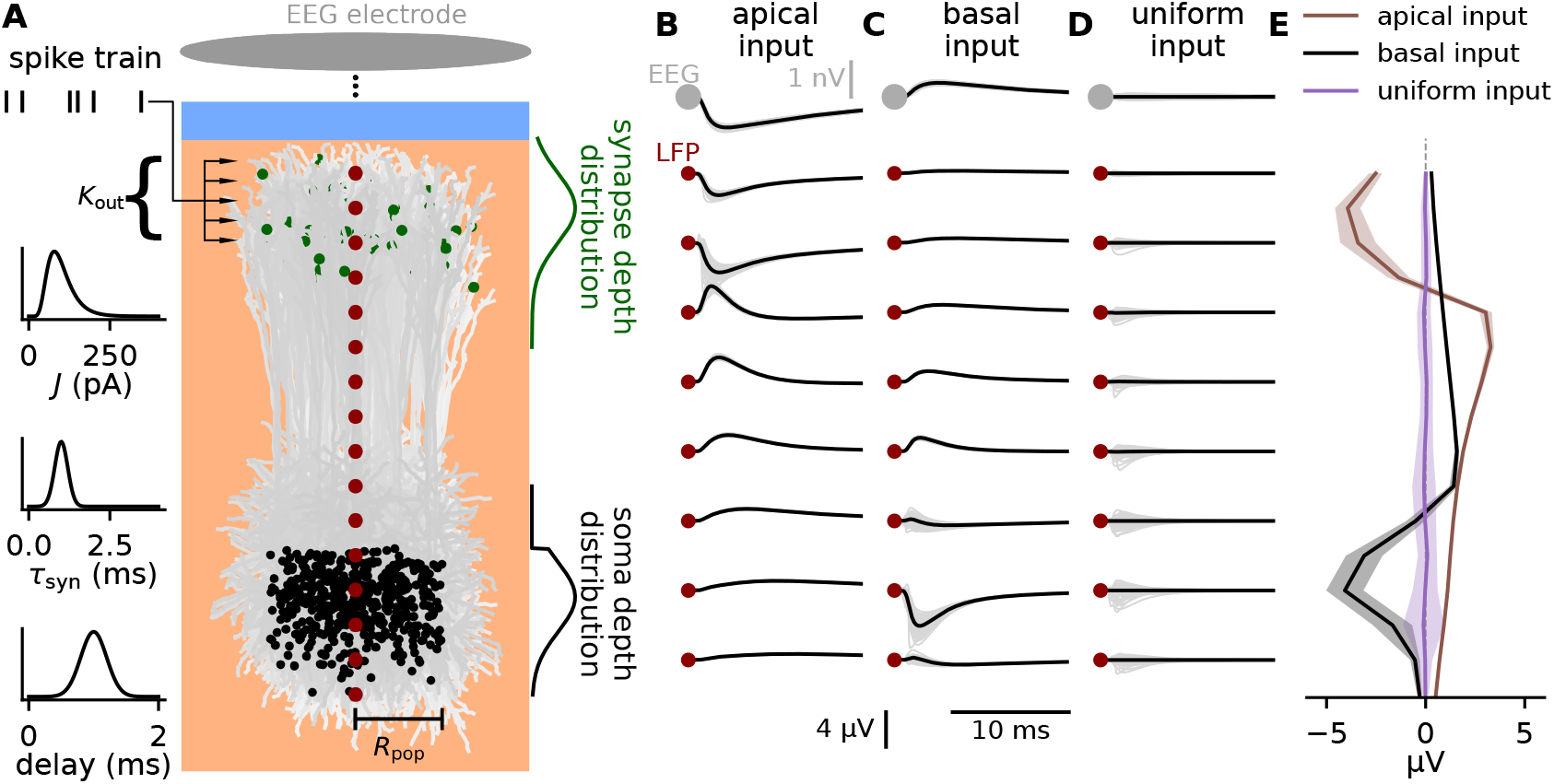
Effect of neuron and synapse heterogeneity on the variability of single-cell LFP kernels. **A:** A population of cortical pyramidal neurons (morphologies depicted in shades of light gray and soma locations as black dots) receives synaptic input from a presynaptic population. Each incoming axon forms, in total, *K*_out_ connections with different postsynaptic neurons. The strength *J* of each synapse is randomly drawn from a lognormal distribution. The synaptic time constant *τ*_syn_ and the synaptic delay are drawn from normal distributions (graphs to the left). The vertical position of each synapse is drawn from a normal distribution (green curve to the right). Some exemplary synapse positions are plotted on the postsynaptic population as green dots. Vertical soma positions are drawn from a capped normal distribution (black curve to the right). Horizontal soma positions are uniformly distributed on a disc within radius *R*_pop_. The LFP response to an activation of all *K*_out_ synapses of a single incoming axon is calculated for different cortical depths (dark red dots). The EEG response outside the head, directly above the population, is calculated using a simple spherical head model. For each parameter configuration, we generate 100 single-cell kernels resulting from different random realizations of neuron and synapse parameters. Each of these kernels describes the postsynaptic LFP (EEG) response to the firing of a different presynaptic neuron. **B–D**: LFP and EEG responses for different synaptic target zones (B: apical; C: basal; D: uniform). Gray: single-cell kernels. Black: population kernel. The “basal input” case is used as the “default case” throughout this study. **E:** Mean (solid curves) and standard deviation (bands) of the maximum LFP deflection at different cortical depths for different synaptic target zones (see legend). See Methods for details on the model and parameter values.

It is well known that the LFP/EEG response of individual cells to synaptic input strongly depends on the location of the synapses [Lindén et al., 2010; Lindén et al., 2011; Næss et al., 2021; Ness et al., 2022]. Since the single-cell kernel is the superposition of such signals from all target cells of a given spike, we expect that this dependence translates into a strong influence of synaptic locations on the shape of the single-cell kernels. Indeed, we find that the single-cell LFP/EEG kernels looked very different when stimulating cells in the population only apically, only basally, or uniformly (Figure 6B-E).

We notice substantial variability in single-cell spike kernels (light gray), however, for the cases of apical or basal input we observe that different single-cell kernels seem to have a similar overall shape, and therefore a pronounced population kernel. In the case of the uniform input, there is more diversity in single-cell LFP kernels, such that the population kernel has very low signal amplitude at all depths. The reason is that individual apical or basal inputs lead to rather stereotypic (but opposite) LFP/EEG responses, irrespective of the exact location of the synapse on the dendrite. In contrast, when considering all possible input locations (uniform) the diversity in the LFP/EEG responses to individual synaptic inputs is larger, leading to substantial cancellation. Furthermore, we notice that the variability seems to be higher close to the input region and decreases with distance from the input region. As a result, there is generally less kernel heterogeneity in the EEG kernels than in the LFP kernels (Figure 6B-D).

By choosing a set of kernels, first from the basal input which we will treat as the “default case” (Figure 6C), and combining them with spike trains (see Methods), we can then calculate the LFP signal by convolving each individual kernel (Figure 7A, gray curves) with its corresponding spike train (Figure 7B, individual spike trains in gray) and summing the results for all single-cell contributions (Figure 7C, gray curves). This is what we treat as ground truth in the following analysis. Further, we convolve the population kernel (Figure 7A, black curves) with the population rate (Figure 7B, black line) to obtain the population kernel LFP (Figure 7C, black curves). For brevity, we first focus on the LFP signal, but the general results also apply to EEG signals, which we will get back to later.

**Figure 7:**
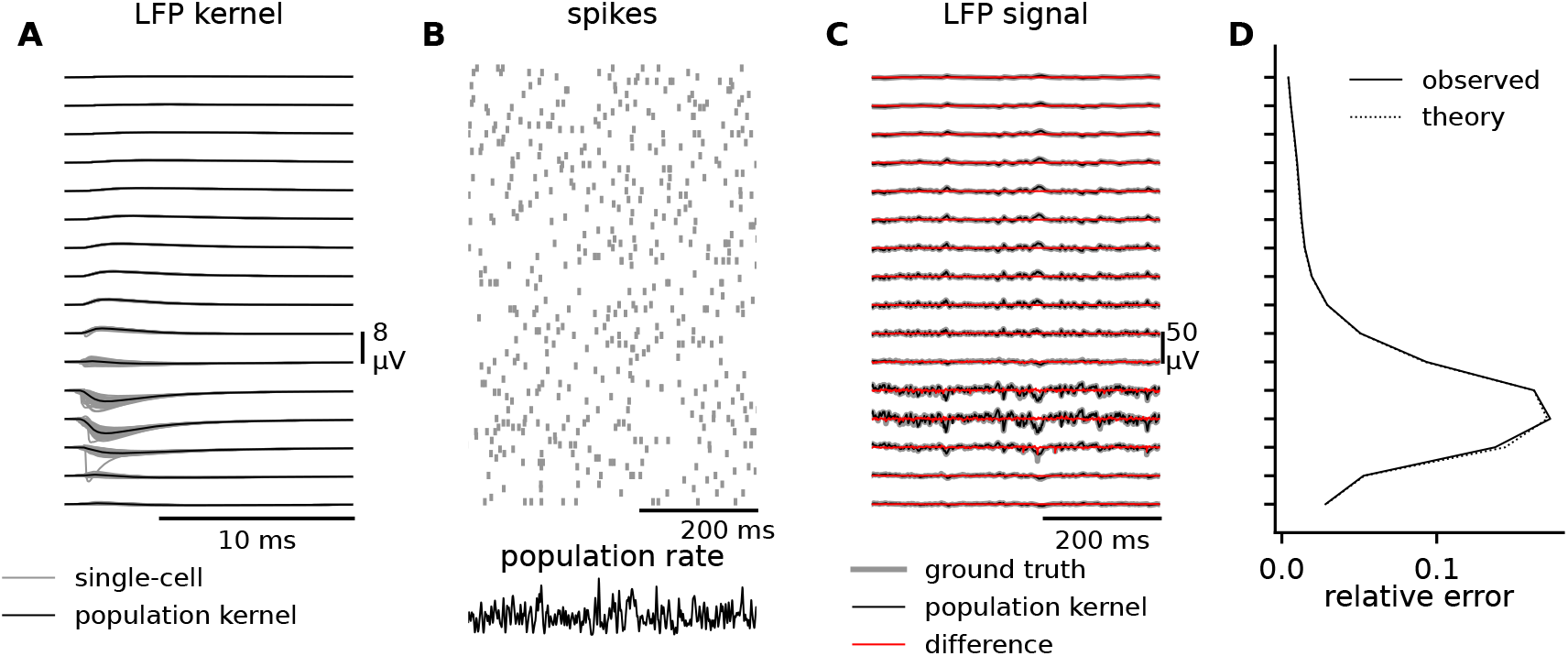
Example of LFP kernels, spike trains, and the resulting LFP signals. **A:** The LFP kernels at different depths (see Figure 6A) with each single-cell kernel in gray and the population kernel in black. The kernels shown here are from the “default” case, corresponding to Figure 6C. **B:** Raster plot of uncorrelated spike trains (see Methods) with a firing rate of 10 s^*−*1^. Below the spikes, the population firing rate (constructed by summing all individual spike trains) is shown in black. **C:** The ground truth LFP signal (gray), the population kernel LFP signal (black), and the difference between them (red), at different depths. **D:** The relative error at different depths (see Methods), either observed from simulations (solid curve) or predicted from theory (dotted curve).

To evaluate the accuracy of the population kernel approach in approximating the ground truth case, we compare the LFP signals (Figure 7C black versus gray curves). We calculate the observed relative error (see Methods), and compare to the relative error predicted from theory, and find them to be almost indistinguishable, demonstrating that the theory is well suited to predict the error (Figure 7D).

We can now evaluate the error of the kernel approach for different parameters of the kernels. To evaluate the relative importance of different factors, we compare different parameter configurations to the “default case” shown in Figure 6C and Figure 7. We start with uncorrelated Poisson spike trains. In the following analysis, we will only show LFP amplitudes and errors but kernels from all tested parameter combinations (see Methods, Table 1) and resulting LFPs are shown in Figure C.13.

**Table 1:** Parameter combinations used for calculating the kernels, where the names of the columns correspond to the parameter combinations tested in Figure 8 and Figure 11. Blank spaces indicate no change from the default values, and only parameters that are varied between simulations are included.

| | default | apical | uniform | small radius | large radius | small $K_{\text{out}}$ | large $K_{\text{out}}$ | similar synapses | variable synapses | narrow input region | broad input region |
| --- | --- | --- | --- | --- | --- | --- | --- | --- | --- | --- | --- |
| $R_{\text{pop}}$ | $250\mu\text{m}$ | | | $125\mu\text{m}$ | $500\mu\text{m}$ | | | | | | |
| $K_{\text{out}}$ | 500 | | | | | 250 | 1000 | | | | |
| syn $z$ SD | $100\mu\text{m}$ | | $10^8\mu\text{m}$ | | | | | | | $50\mu\text{m}$ | $200\mu\text{m}$ |
| syn $z$ mean | $-1270\mu\text{m}$ | $-200\mu\text{m}$ | $-600\mu\text{m}$ | | | | | | | | |
| $\tau_{\text{syn}}$ SD | 0.20ms | | | | | | | 0.10ms | 0.40ms | | |
| syn delay SD | 0.20ms |  |  |  |  |  |  | 0.10ms | 0.40ms |  |  |
| $J$ $s$-value | 0.40nA | | | | | | | 0.20nA | 0.80nA | | |

For basal or apical synaptic input (Figure 8A1, black or brown curves), the ground truth and the population kernel LFP give indistinguishable predictions for the signal amplitude at different depths (the signal amplitude is here represented by the signal standard deviation). This is not the case for the uniformly distributed synaptic input (Figure 8A1, purple curves), which has a much lower amplitude, and a pronounced difference between the ground truth and the population kernel LFP. This is reflected in the error (Figure 8A2) and the relative error (Figure 8A3), where we observe very high relative errors at all depths for the uniform input, and substantially lower error for apical or basal input. Furthermore, for the latter two cases, the error decreases with distance from the input site. This is in agreement with our earlier observations regarding the kernels (Figure 6B-E). Notice also that the observed error (Figure 8A2-A3, solid curves) and the error predicted from theory (Figure 8A2-A3, dotted curves) closely overlap, illustrating again that the theory is perfectly able to predict the error.

**Figure 8:**
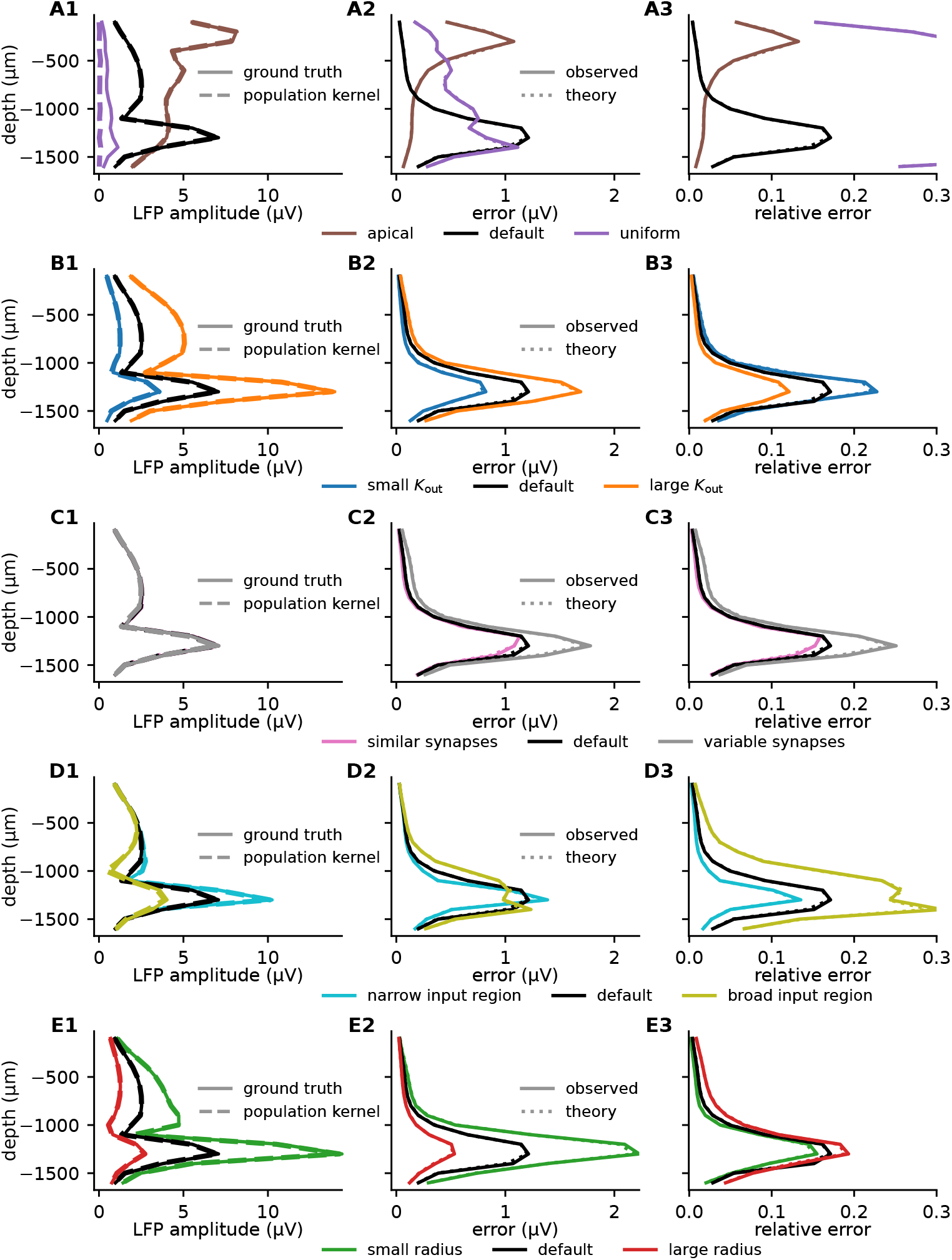
Comparison of how different parameter configurations affect LFP amplitude and population kernel errors. For uncorrelated Poisson input with a rate of 10 s^*−*1^ (see Methods), the figure shows the standard deviation of the LFP at different depths (column 1), and the absolute (column 2) and relative error (column 3) from using the population kernel, for different modifications of the original parameter set (“default”). Each row corresponds to varying a certain feature. **A:** Synaptic input region. **B:** Number of postsynaptic targets *K*_out_ (outdegree). **C:** Variability of synaptic parameters. **D:** Spread of the synaptic input in the depth direction. **E:** Radius of the population.

Intuitively we would expect the number of postsynaptic targets per neuron, *K*_out_, to strongly affect the signal amplitude and the error, since more postsynaptic targets can be expected to increase the amplitude and decrease the variability of the kernels. The reason for this low variability is that each single-cell kernel corresponds to a sum of many extracellular potential responses *h*_*ij*_. These are all “activated” simultaneously by the incoming spike such that differences in *h*_*ij*_ to some degree average out. As a consequence, we would expect the population kernel prediction to become significantly worse if neuronal outdegrees are small. This is indeed the case if we reduce the outdegree *K*_out_ towards lower values (Figure 8B1-B3). A theoretical analysis confirms that the relative error decreases as 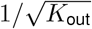 (see Methods).

The synaptic parameters we consider are the synaptic weight, the synaptic time constant, and the synaptic delay. The synaptic weights are lognormally distributed in analogy to Hagen et al. [2016, 2022], while the synaptic time constants and delays are normally distributed. As predicted by the theory (see Methods), decreasing or increasing the standard deviations of these distributions by a factor of two has a negligible effect on signal amplitudes (Figure 8C1), while the error increases with increasing variability (Figure 8C2-C3). We confirmed that the effect on the error is almost entirely determined by the weight distribution, while the time constants and delays have a negligible effect (results not shown).

The spatial spread of the synaptic input is seen to have an important effect on both the signal amplitudes (Figure 8D1) and the errors (Figure 8D2-D3), where a broader region of input gives a much weaker signal and much larger relative errors, similarly to what we saw for the uniformly distributed synaptic input (which can be seen as an extreme case of a broad input region, Figure 6D,E).

When the postsynaptic cells are spatially concentrated, we find a larger LFP amplitude in the center of the population as expected (Figure 8E1). The relative error is however only weakly affected (Figure 8E3).

### 2.4. Sources and effect of spike correlations

To evaluate the error of the kernel approach, we also need to consider the effect of different types of spiking statistics, with different levels of correlation. To this end, we employ the same setup described in the previous subsection but replace the uncorrelated Poissonian input spikes with spike trains generated by two different methods. In a first approach, we create spike trains as realizations of a Multiple Interaction Process (MIP; Kuhn et al. [2003]) with firing rate *v*, fraction *f* of shared spikes, and pairwise correlation coefficient *c* = *f* ^2^. With this model, the firing rate and the level of correlation can easily be controlled, but the auto- and cross-correlations of the resulting spike trains are delta-shaped and thus rather artificial. As an alternative approach, we employ a recurrent point-neuron network model of excitatory and inhibitory neurons (“Brunel network”; [Brunel, 2000]) that can operate in different dynamical regimes and thereby produce spike trains with a more natural correlation structure. Here, we use the same parameters and corresponding network states described in Brunel [2000], and extract spikes from the asynchronous irregular (AI; Figure 9C), and the slow synchronous irregular regime (SI slow; Figure 9D).

**Figure 9:**
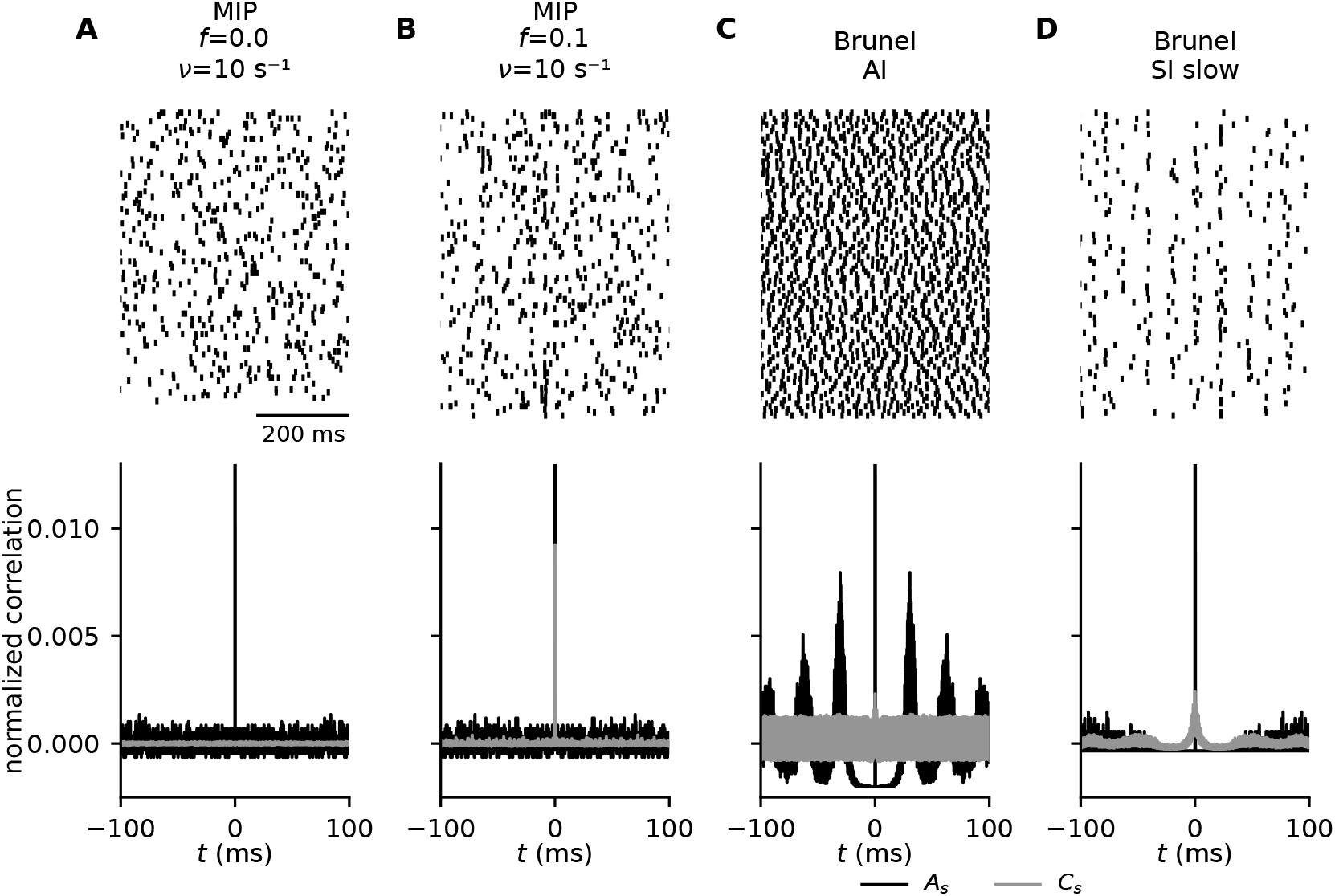
Generating different types of correlated spiking. **A**,**B:** Spiking activity generated by Multiple Interaction Processes (MIP; Kuhn et al. [2003]) with firing rate *v*= 10 s^*−*1^ and correlation coefficients *c* = *f* ^2^ = 0 (A) and 0.01 (B). **C**,**D:** Spiking activity generated by a recurrent network of point neurons [Brunel, 2000] operating in the asynchronous irregular (“AI”; C) and in the slow synchronous irregular regime (“SI slow”; D). Top panels: Raster plots for 100 exemplary neurons. Bottom panels: Normalized spike-train auto- (black) and cross-covariance (gray) functions. The depicted curves represent population averaged correlations obtained from binned spike trains of an ensemble of 100 neurons, with an observation time of 6.1 s, and a binsize of 2^*−*4^ ms. See Methods for details on the spike-generation models and parameter values.

In the parameter configurations discussed above, we used uncorrelated spike trains. However, as earlier discussed, the spike-train correlation will also affect the error (equation (7)). We therefore combine the kernels from the default case used above, with spike trains exhibiting different levels of correlation, including those illustrated in Figure 9. The amplitude of the LFP is highly dependent on the spike trains, and for the MIP spike trains the amplitude increases with both firing rate and correlation (Figure 10A).

**Figure 10:**
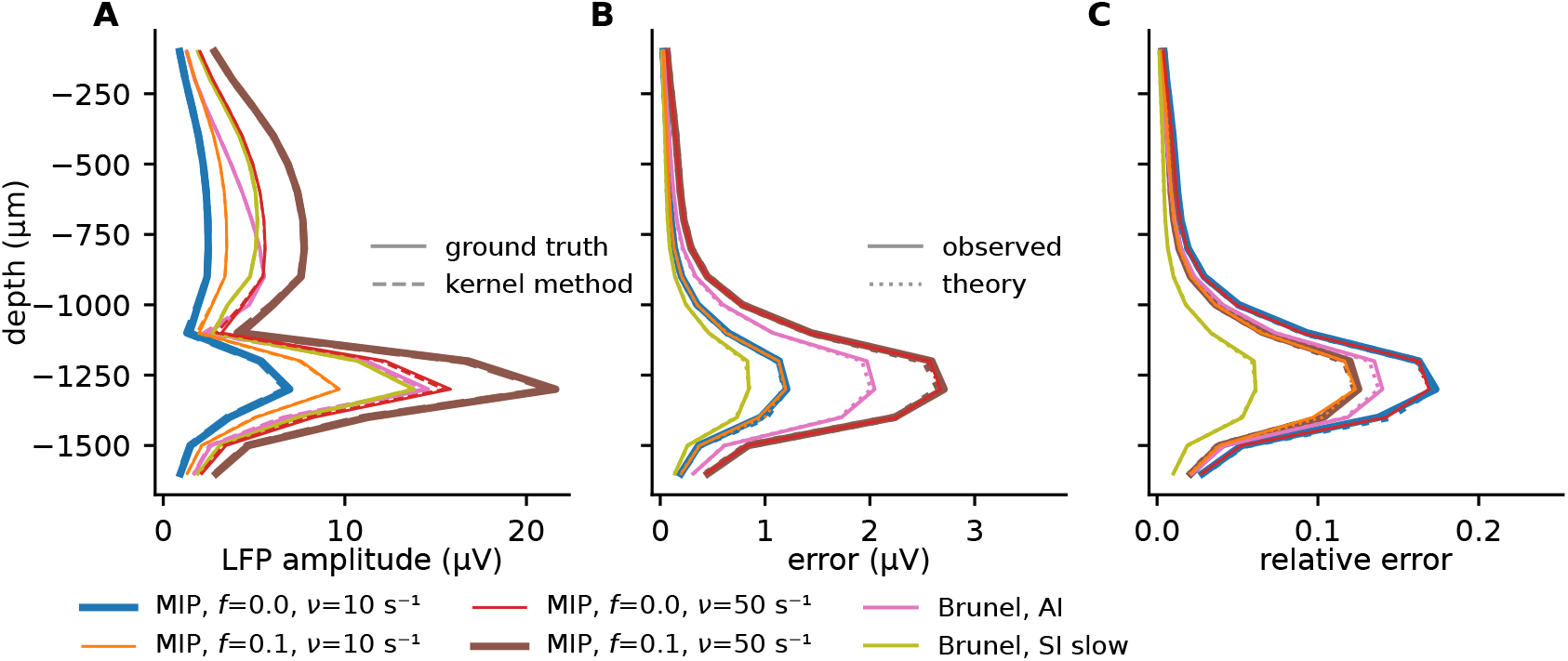
The effect of spiking dynamics in population kernel errors. **A:** For the default kernels (Figure 6C), the amplitude of the LFP signal at different depths is shown for different types of spiking activity, both for the ground truth case (solid lines), and the population-kernel case (dashed lines). **B:** The absolute error across depths, observed in simulations (solid lines) and predicted from theory (dotted lines). **C:** The relative error across depths, observed in simulations (solid lines) and predicted from theory (dotted lines).

The absolute errors from the MIP spike trains appear roughly independent of the correlation, but dependent on the firing rate (Figure 10B), while the relative errors are instead independent of the firing rate but dependent on the correlation. This is confirmed by theory (see Appendix B) and in line with earlier observations in Figure 5B, where we saw in a toy model that the absolute error is only dependent on the correlation for very high levels of correlations (*f >* 0.1). The lowest relative error is from the Brunel SI slow state. This is as expected, because of the highly correlated spiking activity.

### 2.5. Combined effect of kernel heterogeneity and spike-train correlations

We summarize the results in Figure 11A-B, which combines different kernel parameters with different types of spiking activity. If we start by focusing on the kernel parameters (rows), we see that in all cases, uniform synaptic input gives low signal amplitudes and large relative errors. The next highest relative errors are for the case with the broader synaptic input region, which together with the uniform input case demonstrates the importance of the spatial spread of the synaptic input. The lowest relative errors are for the large postsynaptic population (large *K*_out_), followed by the narrow input region. If we instead focus on the different types of spiking activity (columns), we see that the lowest relative error is for Brunel SI slow, while the highest relative error is from the uncorrelated MIP processes.

**Figure 11:**
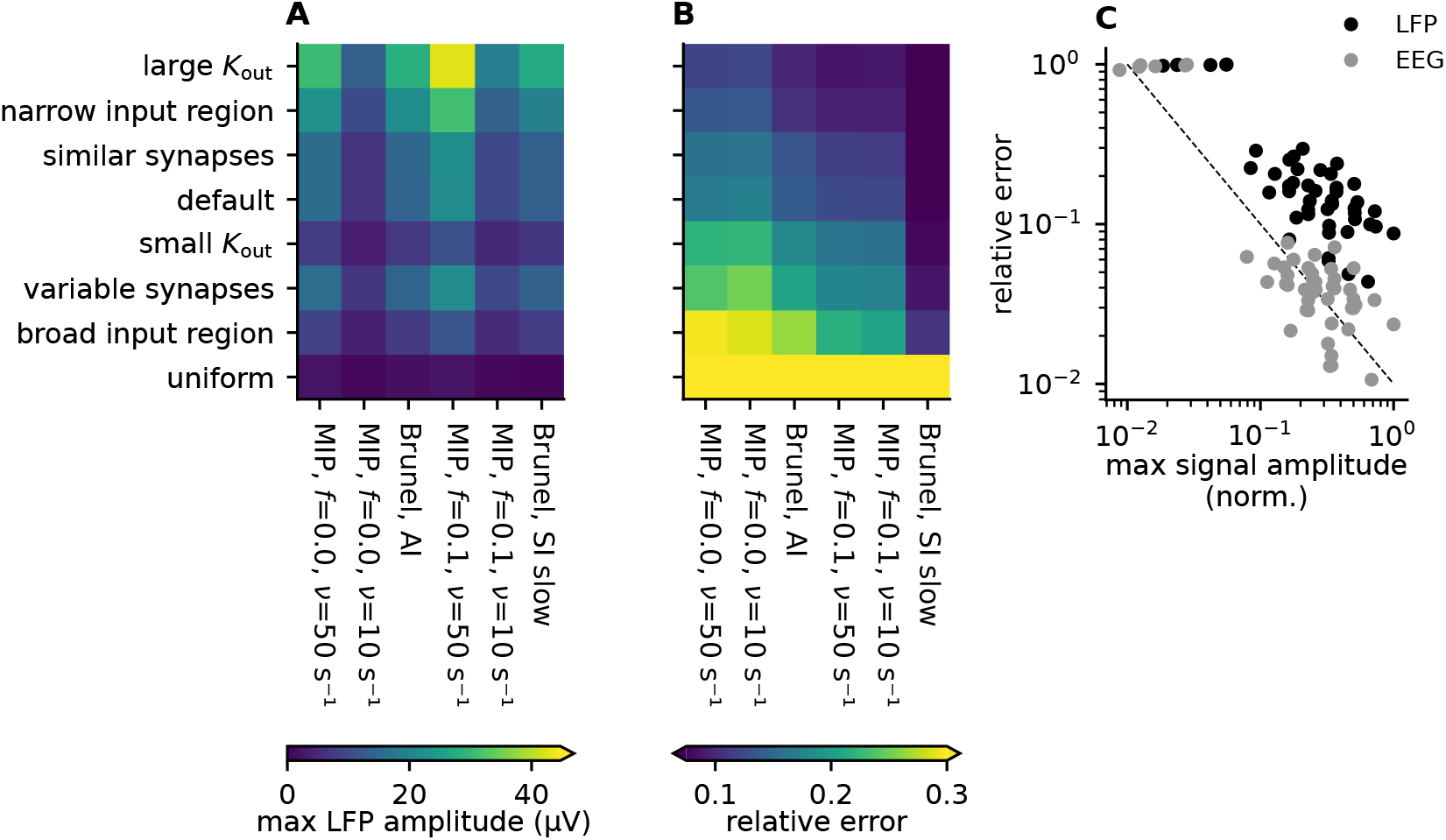
Summary of errors for different kernel parameters and different types of spiking activity. **A:** Maximum ground truth LFP amplitude across depths for different combinations of kernel parameters (rows) and spiking activity (columns). **B:** Maximum observed relative errors across depths, for different combinations of kernel parameters (rows) and spiking activity (columns). The rows and columns are sorted so the largest relative errors are in the bottom left, while the lowest relative errors are in the top right. **C:** The relative error as a function of the signal amplitude for the LFP signal (black dots), and for the EEG signal (gray dots) for all parameter combinations shown in panels A and B. Since the EEG signal intrinsically has a much lower signal amplitude, the LFP and EEG signals are normalized by the maximum observed signal amplitude seen in either of the two signals, so they are easier to visually compare. The dashed line is a visual guideline corresponding to a perfect inverse correlation.

A convenient rule-of-thumb emerges from the results discussed above: The relative error associated with applying the population kernel method is in general inversely proportional to the signal amplitude (Figure 11C). This is an important insight because it means that we can expect the population kernel approach to work best for the synaptic pathways that are dominating the LFP signal, and worst for the synaptic pathways that have a weak LFP contribution. Note that this relationship also holds for the EEG signal, where the error is also substantially lower (Figure 11C, gray dots). As an illustrative example, it has been argued that the LFP and EEG signal is mainly driven by perisomatic inhibitory input to pyramidal cells [Hagen et al., 2016; Teleńczuk et al., 2017; Teleńczuk et al., 2020a,b; Hagen et al., 2022], while excitatory input to pyramidal cells is less important, as it is more uniformly distributed across the postsynaptic pyramidal cells, and therefore gives a relatively weak contribution to the LFP/EEG signal. In this case, we would also expect a large relative error for the excitatory-to-excitatory pathway, but since this synaptic pathway is in this case only associated with a minor LFP/EEG contribution, the high relative error might be acceptable.

## 3. Discussion

### 3.1. Summary

In this study, we have attempted to illustrate what the kernel approach is (Figure 2, Figure 3), and built an intuition for when we can expect it to be applicable (Figure 4, Figure 5). We further developed a mathematical framework to analyze the expected error of the kernel approach and showed that it was capable of accurately predicting the observed errors (Figure 5). From equation (7) we saw that the error was dependent on both the single-cell kernel heterogeneity and the level of correlation between spike trains.

Since LFP, EEG, and MEG signals are, at least in the cortex and in the hippocampus, expected to primarily originate from synaptic input to populations of pyramidal cells, we built a biophysically detailed model population receiving different types of synaptic input, where the individual parameters could be easily adjusted (Figure 6). We then combined these kernels with different types of spiking activity with varying levels of firing rates and correlations (Figure 9). This allowed us to assess how the error introduced by the population kernel approach was affected by different parameter choices for the kernels (Figure 8) and spiking activity (Figure 10).

The results show that the relative error of using the kernel approach will be lowest for the strong signal contributions (e.g., spatially clustered synaptic input and high levels of correlations), and highest for the weak signal contributions (e.g., uniformly distributed synaptic input and low levels of correlations; Figure 11). This implies that those scenarios where the population kernel prediction breaks down are less relevant when considering the total LFP/EEG signal: For cortical scenarios, the LFP/EEG is dominated by apical and basal inputs for which the population kernel prediction only yields a small relative error. Note also that the same holds for LFP signals created by other morphological types of neurons: stellate cells and interneurons lack the asymmetry introduced by the apical dendrites in pyramidal cells. Unless asymmetry is introduced by synapse positions, their LFP contribution can therefore be assumed to resemble the uniform input scenario shown above. The population kernel prediction would break down for populations with symmetric morphologies and synapse distributions. However, their overall contribution to the measured LFP can be expected to be negligible in the presence of pyramidal-neuron LFP contributions.

In summary, these results demonstrate that the kernel approach is a promising method for calculating LFP, EEG, or MEG signals directly from firing rates.

### 3.2. Application to firing rate models

The kernels considered in this paper correspond to the kernels from a single synaptic pathway. Given some prior knowledge or reasonable estimation of synaptic parameters, and how synapses are distributed on postsynaptic neurons, approximate kernels can be derived and used also for firing rate models.

To illustrate its applicability, we here choose a simple population rate model of the form [Montbrió et al., 2015; Schmidt et al., 2018],

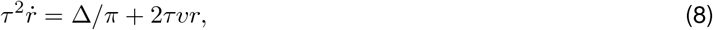

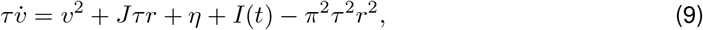

where *r* and *v* are the firing rate and membrane potential, respectively, and *τ* is the membrane time constant. The model is particularly interesting in the context of multi-scale modeling as it has been shown to be an exact macroscopic description of the average dynamics of a population of all-to-all coupled excitatory quadratic integrate-and-fire (QIF) neurons [Montbrió et al., 2015]. The other parameters *J, η* and Δ are derived from the microscopic definition of the QIF network and describe the synaptic weight, and the center and half-width of a Lorentzian distribution of heterogeneous, quenched external inputs, respectively. This population rate model and its dynamic repertoire have been analyzed extensively over the past years with multiple extensions. These include the incorporation of multiple populations to model working memory [Schmidt et al., 2018], inhibitory coupling to produce theta-nested gamma oscillations [Segneri et al., 2020], and sparse coupling and external fluctuations [Goldobin et al., 2021; Di Volo et al., 2022]. The basic model in equation (8) has been shown to produce a non-trivial transient oscillatory behavior upon stimulus-induced (*I*(*t*)) switching between two steady-state attractors (Figure 12A). Using the population kernel prediction such behavior can be modeled in terms of LFP and EEG (Figure 12C), providing the basis for comparisons of population rate model dynamics with experimentally obtained LFPs and EEGs.

**Figure 12:**
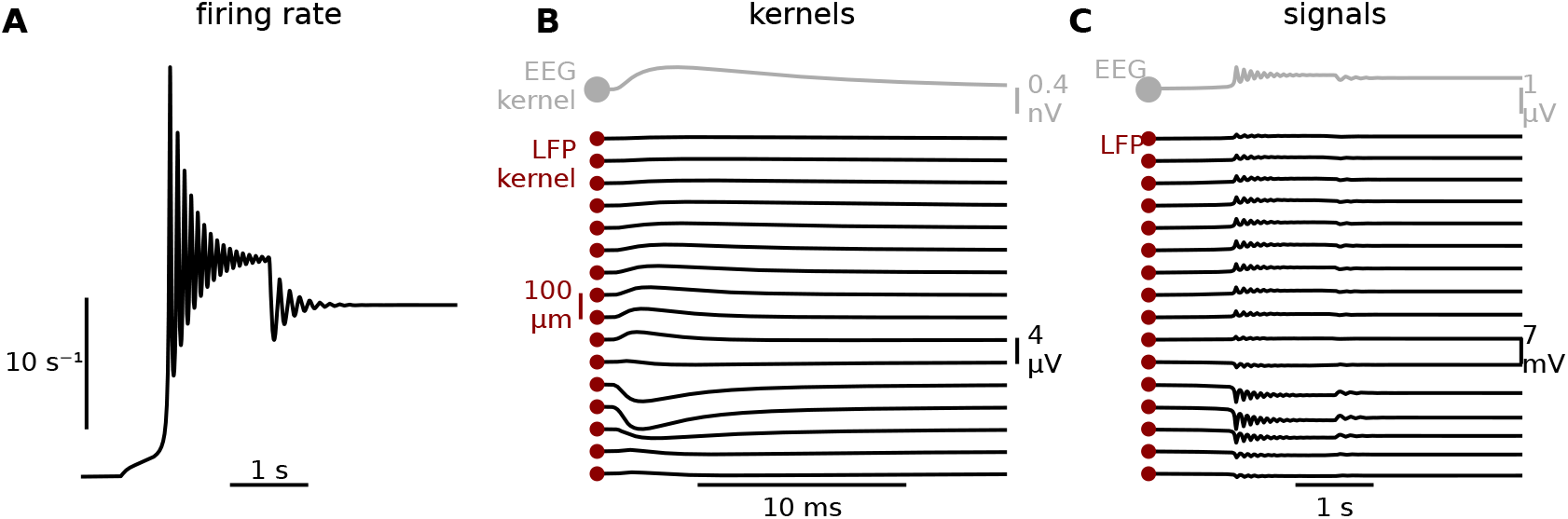
Illustration of the kernel approach applied to a rate model. **A:** Stimulus induced switching dynamics of rate model described by equation (8), with Δ = 2, *η* = *−*10, 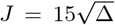, and *τ* = 100 ms [Schmidt et al., 2018]. The stimulus *I*(*t*) is a square pulse with an amplitude of 4, a delay of 1 s, and a duration of 3 s, resulting in switching dynamics similar to what was observed by Montbrió et al. [2015, Fig. 2(a)] **B:** Population kernel for the “default” case (basal input) of the setup introduced in Section “Sources and effect of kernel heterogeneities“ (see Figure 6C). **C:** Transient behavior as observed in the population kernel LFP and EEG signals, calculated by convolving the population rate in panel A with the kernels in panel B. Before the convolution with the LFP/EEG kernel, the population rate is transformed from units of hertz to units of spikes/Δ*t*, and scaled by the considered size of the presynaptic population which was in this case 10,000 [Montbrió et al., 2015].

For firing-rate models with different populations we can combine different kernels for different synaptic pathways. As an example, for an inhibitory-to-excitatory pathway, we could choose a sign-inversed (to change from excitatory to inhibitory input currents) version of the “default” case kernel, to represent perisomatic inhibitory input. Likewise, for an excitatory-to-excitatory pathway, we could use the kernel from uniform synaptic input. All kernels constructed in this study are available online (see Methods), and can in principle easily be modified to accommodate different scenarios.

### 3.3. Limitations

An important caveat of the present study is that we considered a fully linear scenario, with passive postsynaptic neurons and current-based synapses. This allowed us to treat the case, where each single-cell kernel was coupled to its corresponding spike train, as “ground truth”. We could then quantify the error of approximating the LFP/EEG directly from the population kernel and population firing rate. In assuming linearity, we are however ignoring several potentially important factors that may contribute to LFP and EEG signals.

Firstly, we ignored the extracellular action potentials (EAPs) that in principle precede each single-cell kernel. Note that we could in principle easily have included these EAPs in the single-cell kernels by choosing a location for each presynaptic neuron, and calculating the EAP on the recording electrodes from an action potential in the presynaptic neuron. EAPs can have amplitudes of several hundred microvolts if the soma is very close to a recording electrode, but the amplitude falls off rapidly with distance [Pettersen and Einevoll, 2008; Hagen et al., 2015; Halnes et al., 2024], and we would therefore expect a very high single-cell kernel heterogeneity in these EAP contributions. We therefore do not expect that the population kernel would give accurate predictions of EAP-contributions to LFP/EEG signals. However, at least for large cortical populations, we do not expect EAPs to be a major contributor to LFP and EEG signals [Pettersen et al., 2008; Hagen et al., 2022; Ness et al., 2022], but the reader should keep in mind that any putative EAP contribution is neglected in this analysis.

Secondly, in assuming passive postsynaptic neurons, we neglected effects from subthreshold active conductances. It has been demonstrated in modeling studies that subthreshold active conductances can in certain cases be important in shaping the LFP [Ness et al., 2016, 2018], however, this effect can be taken into account also in linear models through linearization [Remme and Rinzel, 2011; Ness et al., 2016, 2018; Hagen et al., 2022]. The effect of other types of non-linearities, such as dendritic action potentials, on the validity of the kernel method LFP estimates should be assessed in future studies.

Thirdly, we relied on current-based instead of conductance-based synapses. Since conductance-based synapses depend on the membrane potential, and change the effective membrane conductance of the postsynaptic neurons, the LFP response to synaptic input will for conductance-based synapses depends on the ongoing synaptic input to the postsynaptic population. It was previously demonstrated by Hagen et al. [2022] that the kernel approach can make accurate LFP predictions also for conductance-based synaptic input, by taking into account the “background level” of synaptic input that each population was receiving. However, while using conductance-based synapses had an important effect on kernel amplitudes [Hagen et al., 2022], it is not expected to have a strong effect on single-cell kernel heterogeneity. Therefore, the error analysis presented here is equally relevant to models using both current-based and conductance-based synapses for calculating kernels.

Also, our analysis here focuses on cortical networks where the LFP/EEG is dominated by inputs onto pyramidal neurons and other contributions are negligible. We show that the relative error of the population kernel method is in general small for large current dipoles, but sizable for overall small signals. It is therefore plausible that the kernel method will work less reliably in other brain regions such as basal ganglia, where there are no pyramidal neurons.

### 3.4. Inference and approximation of population kernels

Population kernels were here constructed from the average of all single-cell kernels for a population of neurons. The latter kernels can be measured in experiments [Swadlow et al., 2002; Bereshpolova et al., 2019; Teleńczuk et al., 2017; Teleńczuk et al., 2020a] and simulations [Hagen et al., 2017; Teleńczuk et al., 2020b], for example using microstimulation of individual neurons. This is, however, experimentally not feasible for a large number of neurons and in simulations it is computationally expensive. Since variability in single-cell LFP kernels is low in some scenarios, we can expect that an approximation of the population kernel based on single-cell kernels of a small subpopulation is still valid, and indeed Hagen et al. [2022] demonstrated that population kernels could be accurately estimated based on a single biophysically detailed cell simulation.

A direct way to obtain population kernels is via simultaneous stimulation of the whole population of neurons, for example using optogenetic techniques. Also, population kernels can be inferred via deconvolution techniques [Mukamel et al., 2005] from given compound extracellular potentials and population rates. This procedure, however, relies on the fact that those neurons from which spike trains are recorded are those with the dominant single-cell kernels. If other populations of neurons from which no spikes are recorded contribute significantly to the extracellular potential, then the inferred population kernel is invalid. In the case of spike recordings from multiple populations, one can use the MIMO (multiple input - multiple output [Perreault et al., 1999]) scheme for deconvolutions of the individual population kernels.

### 3.5. Definition of population

Typically a population is defined via common input statistics and physiological parameters between neurons such that output spiking statistics are similar. Here, we need in addition that the single-cell spike kernels of neurons in a population are similar. This includes similar postsynaptic targets and projection patterns to them as well as passive properties of postsynaptic targets. So the definition of a population is not only based on incoming connection statistics but also on outgoing connection statistics. Also what defines a population might dynamically change: if correlations, i.e., spiking statistics, between two populations are large then merging them into one population even if they have different kernels would lead to a good population-kernel prediction.

### 3.6. Conclusion

As reviewed in the Introduction, several different approaches to calculate LFP/EEG/MEG signals from point-neuron or firing rate models have been suggested [Deco et al., 2008; Sanz-Leon et al., 2015; Mazzoni et al., 2015; Teleńczuk et al., 2020a; Glomb et al., 2022; Tesler et al., 2022], but quantitative evaluations of the accuracy of these approaches have often been lacking. Here, we have presented a thorough analysis of how the kernel method works, and when we can expect it to be a good approximation. Our results further establish the kernel approach as a promising method for calculating brain signals from large-scale neural simulations, and we hope that the kernel approach can therefore be used with more confidence.

## 4. Methods

### 4.1. Forward modeling

The calculation of the extracellular potential was done using a well-established forward-modeling scheme based on electrostatics with current sources computed via solving the membrane potential dynamics of each cell given all its inputs [Halnes et al., 2024]. For the neural simulations we used LFPy [Hagen et al., 2018], running on top of NEURON [Carnevale and Hines, 2006].

#### 4.1.1. Calculating EEG signals

Current dipole kernels were calculated from the neural simulations using LFPy [Hagen et al., 2018], and could in principle be used with arbitrarily simple or detailed head models. EEG signals were calculated with the four-sphere head model [Næss et al., 2017] implemented in LFPy.

#### 4.1.2. Toy model for spike-LFP kernel

The spike-LFP kernels from the toy model (Figure 3, Figure 4, Figure 5) were double exponential functions (rise time *τ*_1_ = 0.2 ms, decay time *τ*_2_ = 1 ms), which only varied in amplitude *A*_*i*_. The implementation was equivalent to the “Exp2Syn” mechanism in NEURON, and given by,

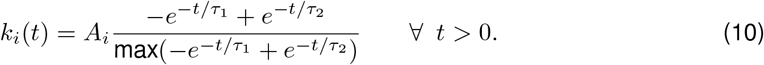

The mean amplitude was always 1.0 V, while the standard deviation of the amplitude was varied as detailed in the individual figures. The time resolution of the simulations was 0.1ms.

### 4.3. Biophysically detailed simulations

We used the rat cortical layer 5 pyramidal cell model from Hay et al. [2011], where all active conductances were removed to make the cell passive [Ness et al., 2016, 2018]. We used current-based synaptic input with exponential decay, and a time resolution of 2^*−*4^ms. For calculating single-cell spike-LFP kernels, we generated for each presynaptic neuron *j*, a population of *K*_out_ (default value: 500) postsynaptic instances of the pyramidal cell model. The cells were aligned with and randomly rotated around the *z*-axis, and the *z*-positions of the somas were drawn from a capped normal distribution (mean: −1270*µ*m, SD: 100*µ*m). The cap was introduced to avoid neurons protruding out of the cortex. The somas were uniformly distributed in the *xy*-plane within a radius *R*_pop_ (default value: 250*µ*m). Each postsynaptic neuron *i* had a single synapse with weight *J*_*ij*_ drawn from a lognormal distribution, calculated through `scipy.stats.lognorm` (mean: 0.1nA, default *s*-value: 0.4nA, see `scipy.stats.lognorm` documentation). The spatial distribution of the synapses in the depth direction was drawn from a normal distribution (default syn z mean: −1270*µ*m, default syn z SD: 100*µ*m). The synaptic time constants *τ*_syn_ were drawn from a normal distribution (mean: 1ms, default SD: 0.2ms), as were synaptic delays (syn delay mean: 1ms, default syn delay SD: 0.2ms). The default values of the parameters as well as the different variations tested in this study are listed in Table 1.

### 4.4. Synthetic spike-trains with correlations

Synthetic spike trains with varying levels of correlations and firing rates were generated through Multiple Interaction Processes (MIP) [Kuhn et al., 2003]. Here, a “mother spike train” was first generated with the same firing rate as the target spike trains. The spike times were generated through a homogeneous Poisson process using Elephant [Denker et al., 2018]. For each “child spike train”, a fraction *f* of the spikes where randomly selected from the mother spike train, while the remaining spikes were generated through homogeneous Poisson processes. Consequently, *f* varies between 0 and 1, and *f* = 0 corresponds to uncorrelated homogeneous Poisson processes, while *f* = 1 corresponds to fully correlated (identical) spike trains. Since each spike is copied with probability *f* ^2^ into two child spike trains, the correlation coefficient of the latter is given by *c* = *f* ^2^ (for details see Appendix B).

### 4.5. Error measures

We define the absolute squared error of the population kernel signal 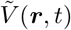 in approximating the ground truth signal 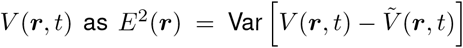, where Var […] denotes the variance across time, computed separately for each electrode position ***r***. The relative squared error is defined as 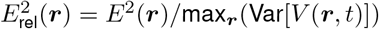, that is, the absolute squared error at each electrode, normalized by the maximum (over the electrodes) variance of the ground truth signal. Note that the error is normalized by the largest value of the ground truth signal variance because the ground truth signal will often have electrodes with very near-zero signal amplitudes, and therefore very high, but irrelevant relative errors. In the case of the toy model (Equation (10)) that is agnostic to spatial positions, the normalization does not involve a maximum over the electrodes.

### 4.6. Firing rates

Firing rates are constructed from spike trains by counting the number of spikes within time steps of length Δ*t*, and normalizing by Δ*t*. For kernels generated with the toy model (Figures 3-5), the time step duration is Δ*t* = 0.1 ms, while for the simulations with biophysically detailed kernels (Figures 6-12), it is Δ*t* = 2^*−*4^ ms.

### 4.7. Point-neuron network simulation

The point-neuron network model was a random balanced network with delta synapses [Brunel, 2000], based on the `brunel_delta_nest.py` example that comes with NEST. We used NEST 3.6 [Villamar et al., 2023], with the same network parameters and network states as Brunel [2000], that is, we extracted spikes from an asynchronous irregular (AI) regime (*g* = 5, *η* = 2.0, *J* = 0.1) [Brunel, 2000][Figure 8C], and a slow synchronous irregular (SI) regime (*g* = 4.5, *η* = 0.9, *J* = 0.1) [Brunel, 2000][Figure 8D]. The time resolution was 0.1 ms.

### 4.8. Mathematical derivation of error estimate

In simulations, we measure the squared error

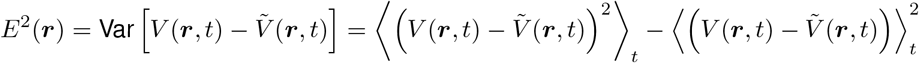

of the population kernel method as the variance of the difference signal 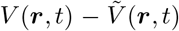 across time. By definition, due to the time average ⟨·⟩, the error depends on the statistics of spike trains *s*. In addition, in principle, it depends on all details of the single-cell spike-LFP kernels *k*. Yet, for networks of biologically realistic size, the LFP is made up of many contributions, such that the squared error *E*^2^(***r***) will not vary too much between different statistically equivalent realizations of single-cell kernels. Therefore, the expectation ⟨*E*^2^(***r***)⟩_*k*_ across different realizations of single-cell kernels *k* can be assumed to be informative about the error *E*^2^(***r***) for one particular realization.

The expected squared error is

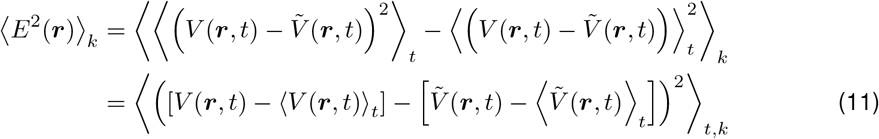

Inserting the definition of the ground truth LFP (equation (3)) and the population-kernel approximation (equation (6)) yields the error expression (equation (7)) of the main text (for details see Appendix A).

For the prediction of the relative squared error, we divide the error by the variance of the ground truth LFP

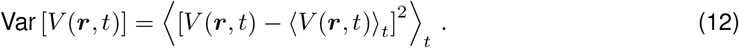

On expectation, the latter time average can be calculated analogously to the error (for details see Appendix A). This allows us to obtain some intuition on the expected relative squared error

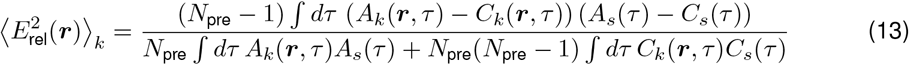

in relation to features of single-cell kernels. To do so, we employ equation (4) and write the single-cell kernel in terms of individual extracellular potential responses *h*_*ij*_(***r***, *t*) = *ξ*_*ij*_*J*_*ij*_*χ*_*ij*_(***r***, *t*), where we explicitly split the synaptic strength *J*_*ij*_ and the adjacency values *ξ*_*ij*_ ∈{0, 1} from the impulse response *χ*_*ij*_(***r***, *t*). The latter characterizes the LFP response, measured at time *t* and location ***r***, to a unit input arriving at the synapse between neurons *i* and *j*. The single-cell spike-LFP correlations then read

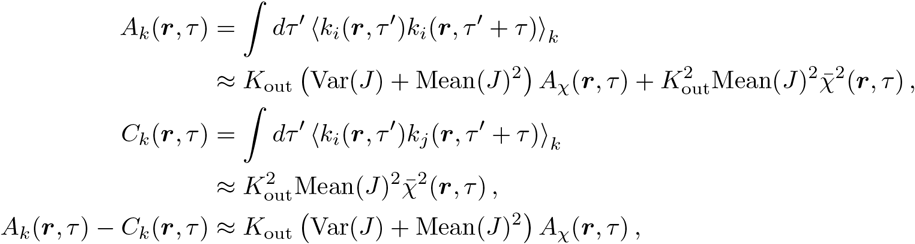

with Mean(*J*) and Var(*J*) denoting the mean and variance of synaptic weights, impulse-response statistics 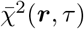 and *A*_*χ*_(***r***, *τ*), and *K*_out_ the outdegree of presynaptic neurons (see Appendix A). Interestingly, the error 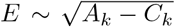 (square root of numerator in equation (13)) - due to cancellations between *A*_*k*_ and *C*_*k*_ - scales as 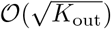 (Figure 8B2), while the signal standard deviation 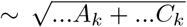 (square root of denominator in equation (13)) scales as 𝒪(*K*_out_) (Figure 8B1), such that the relative error decreases with postsynaptic population size as 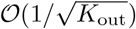 (Figure 8B3). Furthermore, the signal standard deviation is roughly independent of the variability in synaptic strengths (Figure 8C1). This variability Var(*J*) only enters in the term of *A*_*k*_ that is proportional to *K*_out_ and thus subleading compared to the other terms in *A*_*k*_ and *C*_*k*_ that are proportional to 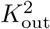. In the error these terms proportional to 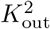 exactly cancel, such that the error increases with larger variability in synaptic strengths (Figure 8C2). The closer the different synaptic locations *k* and *l* are (see narrow vs broad input region or apical/default vs uniform), the larger the product of different impulse responses *χ*_*ki*_(***r***, *τ* ^*′*^)*χ*_*li*_(***r***, *τ* ^*′*^+ *τ*). Therefore, the signal standard deviation, which contains products of different impulse responses in 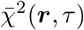 (see Appendix A), grows when synaptic locations become more similar (Figure 8A1,D1). In contrast, the error only depends on *A*_*χ*_(***r***, *τ*), which in turn only depends on products of the same impulse responses (see Appendix A). Therefore, the error is less sensitive to the width of the input region (Figure 8D2). Still, both impulse response statistics 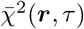 and *A*_*χ*_(***r***, *τ*) depend strongly on the type of input region, leading to a strong dependence of the signal standard deviation and error on the input region (Figure 8A2). Also, both terms increase the smaller the radius of the population, because LFP-generating sources are closer to the recording electrode. Therefore, both the signal standard deviation (Figure 8E1) and the error (Figure 8E2) increase with smaller population radius.

### 4.9. Code availability

Simulation code to reproduce all figures in this paper, as well as all the simulated kernels are freely available from https://github.com/torbjone/kernel_validity.git.

## Acknowledgements

This work received funding from the European Union Horizon 2020 Research and Innovation Programme under Grant Agreement No. 945539 [Human Brain Project (HBP) SGA3], No. 101147319 [EBRAINS 2.0], the Norwegian Research Council (NFR) through NOTUR (No. NN4661K), the Helmholtz Association Initiative and Networking Fund (SO-092, Advanced Computing Architectures), and the Helmholtz Metadata Collaboration (HMC; ZT-I-PF-3-026). Open-access publication is funded by the Deutsche Forschungsgemeinschaft (DFG, German Research Foundation) - 491111487. We are grateful to our colleagues in the NEST developer community for continuous collaboration. All network simulations were carried out with NEST (http://www.nest-simulator.org).

## Appendix A. Derivation of error formula

Here we provide details for the derivation of the expected squared error between the ground truth LFP and the population kernel approximation. Inserting the definitions

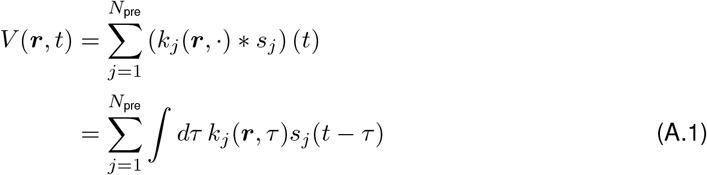

and

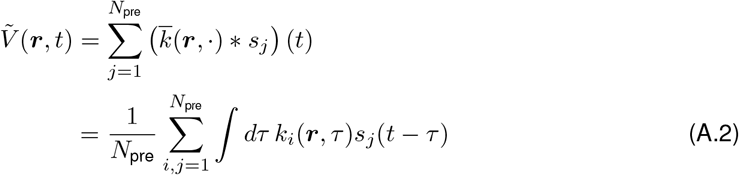

into equation (11), we obtain

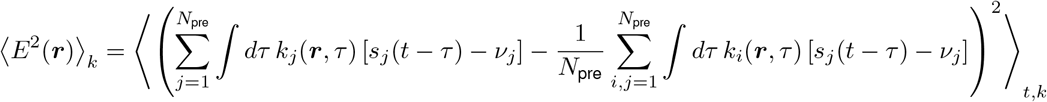

with firing rates *v*_*j*_ = ⟨*s*_*j*_(*t* − *τ*)⟩_*t*_. Multiplying out the square then yields

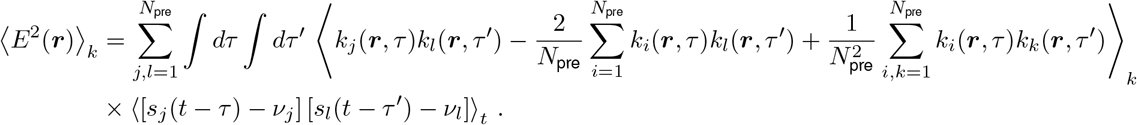

The averages over time ⟨·⟩_*t*_ yield the spike-train covariances *c*_*jl*_(*τ* ^*′*^−*τ*) = ⟨[*s*_*j*_ (*t* − *τ*) − *v*_*j*_] [*s*_*l*_(*t* − *τ* ^*′*^) − *v*_*l*_]⟩ _*t*_, which for stationary spike-train statistics only depends on the relative time between spike trains. For the average over single-cell spike-LFP kernels ⟨·⟩_*k*_ one splits the sum ∑ into a sum over equal indices ∑_*j*_ and a sum over unequal indices ∑_*j*≠*l*_ to obtain after some simplifications

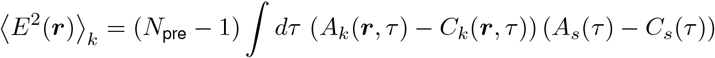

with population-averaged spike-train autocovariance 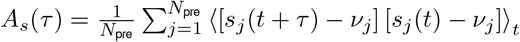, population-averaged spike-train cross-covariance 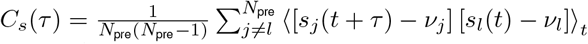, single-cell spike-LFP kernel autocorrelation *A*_*k*_(***r***, *τ*) = ∫*dτ* ^*′*^ ⟨*k*_*i*_(***r***, *τ* ^*′*^)*k*_*i*_(***r***, *τ* ^*′*^ + *τ*)⟩_*k*_, and single-cell spike-LFP kernel cross-correlation *C*_*k*_(***r***, *τ*) = ∫*dτ* ^*′*^ ⟨*k*_*i*_(***r***, *τ* ^*′*^)*k*_*j*_(***r***, *τ* ^*′*^ + *τ*)⟩_*k*_ for *i* ≠ *j*. In practice, to calculate *A*_*k*_ and *C*_*k*_, one replaces the expectation value over single-cell spike-LFP kernel statistics by an empirical average that can be measured

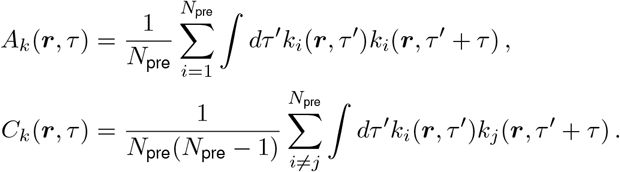

Analogous to the calculation above, the variance of the ground truth LFP can be calculated on expectation

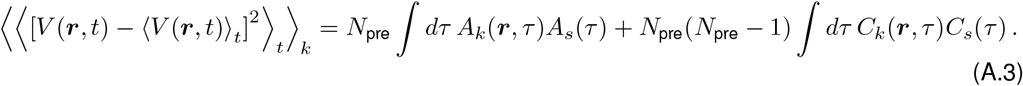

By expressing *A*_*k*_ and *C*_*k*_ in terms of impulse responses (equation (4)), we obtain

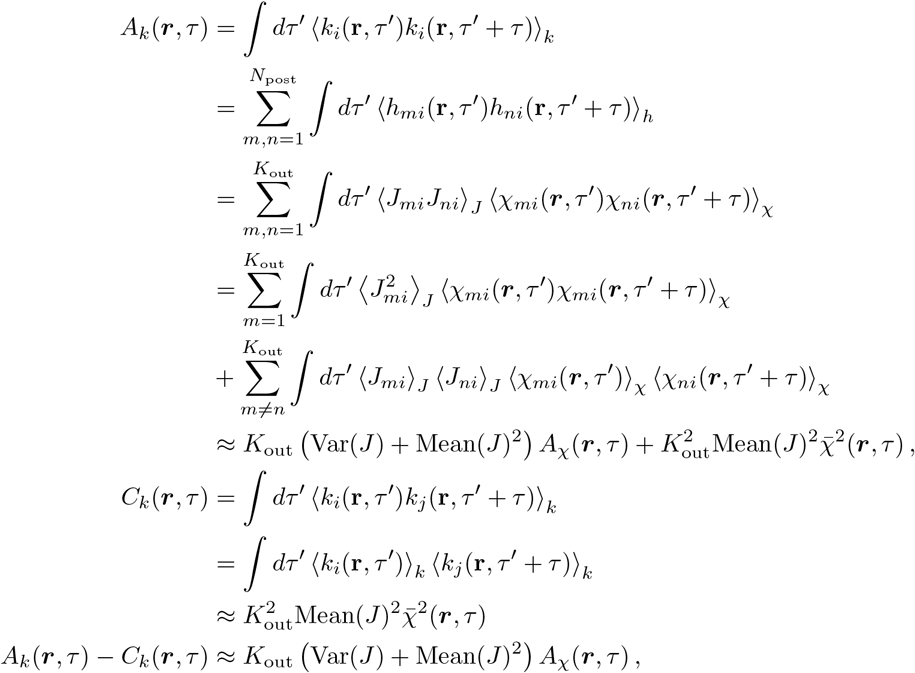

with Mean(*J*) = ⟨*J*_*ki*_⟩_*J*_ and Var 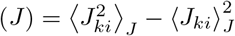 the mean and variance of synaptic weights, *K*_out_ the outdegree of presynaptic neurons, and impulse-response statistics

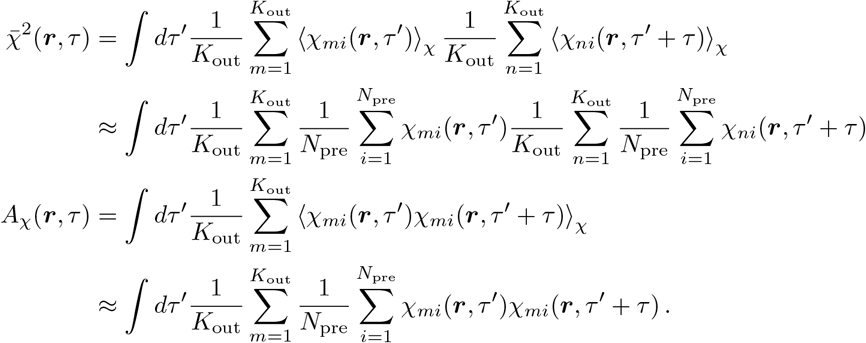

## Appendix B. MIP spike train generation and correlations

Let’s consider a homogenous Poisson spike train *m*(*t*) (“mother spike train”) of rate *v*und independent Poisson spike trains *S*_*i*_(*t*) of rate (1− *f*)*v*. We define child spike trains *s*_*i*_(*t*) as a superposition of *S*_*i*_(*t*) and *m*_*i*_(*t*), where *m*_*i*_(*t*) is a Poisson process of rate *fv* that consists of a randomly chosen fraction *f* of spikes from the mother spike train *m*(*t*). By definition, each child spike train is then a Poisson process with rate *v*and auto-covariance *A*_*s*_(*τ*) = *vδ*(*τ*). Since each spike of the mother spike train is selected with probability *f* ^2^ to be copied into *m*_*i*_(*t*) and *m*_*j*_(*t*), the child spike trains *s*_*i*_ and *s*_*j*_ share a common Poisson spike train of rate *f* ^2^*v*. The cross-covariance between child spike trains is therefore *C*_*s*_(*τ*) = *f* ^2^*vδ*(*τ*), and the correlation coefficient is *c* = *f* ^2^.

Since both auto- and cross-covariances of MIP spike trains are proportional to the firing rate, the latter exactly cancels in the relative error of the population kernel approximation (equation (13)). The absolute error is given by the difference *A*_*s*_(*τ*) − *C*_*s*_(*τ*) = *v*(1 −*c*)*δ*(*τ*) and therefore rather insensitive to correlations *c* that are small (Figure 5B, Figure 10B). For the signal amplitude, cross-covariances are, however, amplified by a factor *N*_pre_ (equation (A.3)), leading to a strong dependence on *c* of the signal amplitude (Figure 5A, Figure 10A) and the relative error (Figure 5C, Figure 10C).

## Appendix C. All kernels

**Figure C.13:**
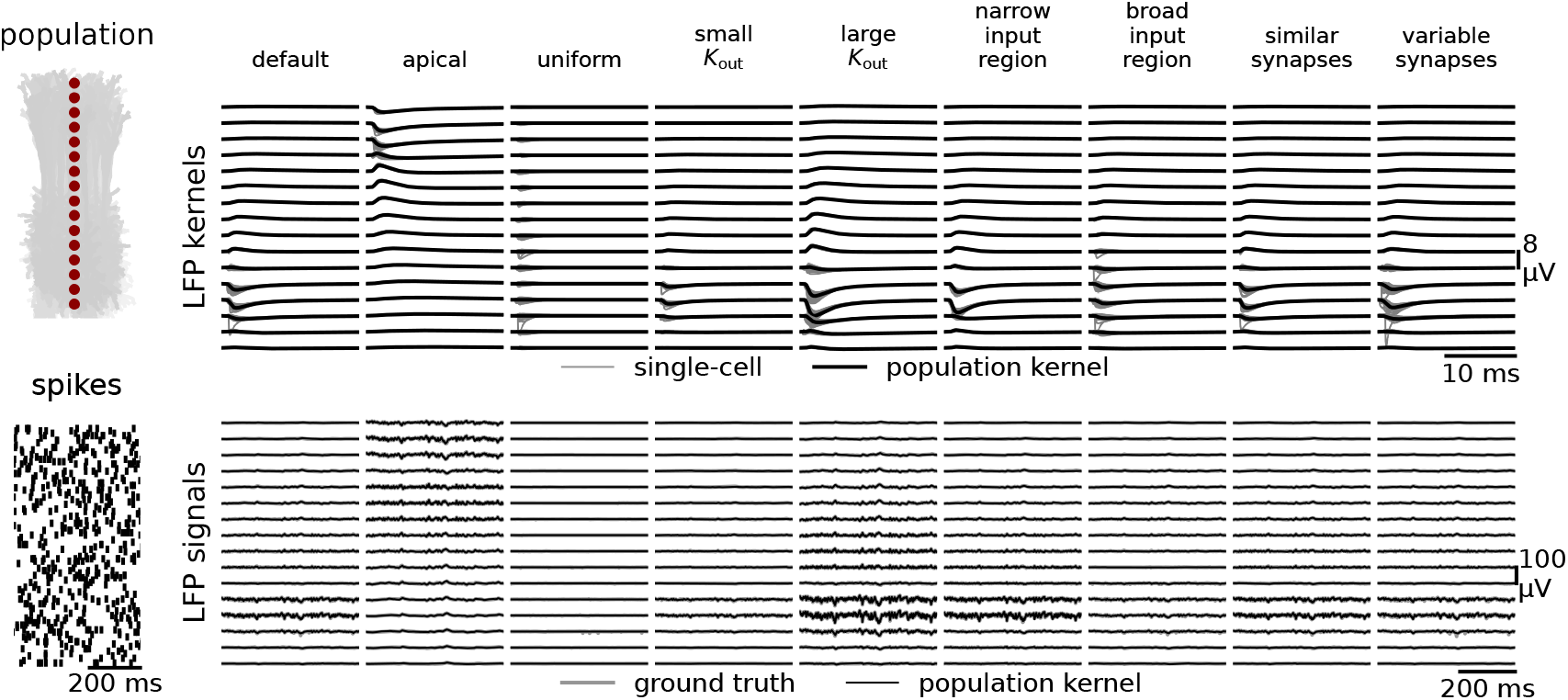
All tested LFP kernels and the resulting LFP signals. The LFP kernels for all tested parameter combinations in Table 1 (top row), and the resulting LFP signals (bottom row). The spike trains were from an uncorrelated Poisson process with a firing rate of 10 s^*−*1^.

